# Cleavage cascade of the sigma regulator FecR orchestrates TonB-dependent signal transduction

**DOI:** 10.1101/2025.01.06.631622

**Authors:** Tatsuhiko Yokoyama, Ryoji Miyazaki, Takehiro Suzuki, Naoshi Dohmae, Hiroki Nagai, Tomoya Tsukazaki, Tomoko Kubori, Yoshinori Akiyama

**Affiliations:** Department of Microbiology, Graduate School of Medicine, Gifu University, 1-1 Yanagito, Gifu 501-1194, Japan; Division of Biological Science, Graduate School of Science and Technology, Nara Institute of Science and Technology, 8916-5 Takayama, Ikoma, Nara 630-0192, Japan; Biomolecular Characterization Unit, RIKEN Center for Sustainable Resource Science, 2-1 Hirosawa, Wako, Saitama 351-0198, Japan; Center for One Medicine Innovative Translational Research (COMIT), Institute for Advanced Study, Gifu University, 1-1 Yanagito, Gifu 501-1194, Japan; Institute for Life and Medical Sciences, Kyoto University, 53 Shogoinkawahara-cho, Sakyo-ku, Kyoto 606-8507, Japan

**Keywords:** iron transport, Fec system, Ton system, twin-arginine translocation system, signal transduction

## Abstract

TonB-dependent signal transduction is a versatile mechanism observed in gram-negative bacteria, integrating energy-dependent substrate transport with signal relay. In *Escherichia coli*, the TonB-ExbBD motor complex energizes the TonB-dependent transporter FecA, facilitating ferric citrate import. FecA also functions as a sensor, transmitting signals to the cytoplasmic membrane protein FecR. We previously demonstrated that FecR undergoes a three-step cleavage process, culminating in the activation of the cytoplasmic sigma factor FecI, which drives *fec* gene transcription. Here, we describe the complete mechanism of FecR cleavage-mediated ferric citrate signaling involving FecA and TonB. The cleavage cascade begins with FecR autoproteolysis prior to membrane integration. The soluble C-terminal domain (CTD) fragment of FecR is co-translocated with the N-terminal domain (NTD) fragment through a Tat system-mediated process. In the periplasm, the interaction between the CTD and NTD fragments prevents further cleavage. This inhibition is lifted by TonB-mediated motor function, which releases the CTD, allowing the cleavage cascade to proceed. This process is essential for ferric citrate signal-induced activation of *fec* gene expression. Our findings reveal that the regulation of FecR cleavage, relying on the TonB-FecA axis, plays a central role in bacterial response to ferric citrate signals.

**Significance Statement:** Scarcity of iron, an essential element for life, has driven bacteria to evolve intricate acquisition systems, yet the molecular basis of their signal transduction mechanisms remains elusive. Unlike conventional pathways, iron transport systems employ outer membrane receptors that mediate signal transduction across both the outer and cytoplasmic membranes, powered by TonB. Using the *Escherichia coli* Fec system, we uncovered the mechanism by which FecR’s cleavage cascade orchestrates ferric citrate signaling through TonB-mediated energy transfer. These findings shed light on similar mechanisms in gram-negative bacteria and have significant implications for understanding bacterial adaptation and pathogenesis.

## Introduction

Bacteria adapt to changing environmental conditions by sensing external molecules and transducing signals across their membranes. In many bacterial signal transduction systems, inducers are transported into the cytoplasm to initiate regulatory responses, as demonstrated by the lactose operon regulation (1) and quorum sensing systems (2). In some gram-negative bacteria, outer membrane proteins detect ligands and relay signals to interacting partners, including inner membrane or soluble proteins, ultimately influencing cytoplasmic gene expression (3). Iron transport mechanisms, extensively studied in this context, reveal critical insights into iron acquisition pathways and their associated regulatory processes (4). Notably, in the Fec transport system of *Escherichia coli*, the ligand ferric citrate is not transported into the cell; only iron is imported (5), highlighting the pivotal role of periplasmic signaling relays.

The Fec transport system comprises the outer membrane protein FecA (6, 7), the periplasmic protein FecB, and the inner membrane complex FecCDE, an ABC transporter (8). These components are encoded by the *fecABCDE* operon (*fec* operon), whose transcription is regulated by FecI, an alternative sigma factor (9, 10). FecA serves as both a transporter and sensor for external ferric citrate, transmitting signals to FecI to activate *fec* operon expression (11). A key step in this pathway is the periplasmic interaction between FecA and FecR, a type II (N_IN_-C_OUT_) cytoplasmic membrane protein involved in signal relay (12–16).

Studies from the 1970s established the essential role of TonB in chelate-mediated iron uptake, acting after signal reception (17–19). Further research using the FhuA transporter demonstrated that TonB-dependent iron transport is energy-driven, powered by the Ton system (20). The Ton system functions as a molecular motor, composed of TonB, ExbB, and ExbD. ExbB and ExbD form a proton channel across the cytoplasmic membrane, utilizing the proton motive force (PMF) to generate mechanical energy for TonB activity, which is hypothesized to involve either rotational or pulling motions (20, 21). TonB’s C-terminal periplasmic domain interacts with the TonB box of outer membrane transporters, inducing conformational changes in TonB-dependent transporters (TBDTs) to facilitate substrate transport (20–22).

This study investigated the signaling mechanism of the Fec system in the context of TonB function. Previously, we demonstrated that (i) FecR undergoes three sequential cleavage steps to produce distinct N-terminal fragments, (ii) ferric citrate signal induces the second cleavage, and (iii) the S2P family intramembrane protease RseP catalyzes the final cleavage, releasing a fragment that associates with FecI to activate *fec* operon transcription (**Fig. 1A**) (23). Herein, we elucidate how TonB-mediated mechanical energy is harnessed by FecA to trigger the FecR cleavage cascade, enabling downstream gene regulation.

**Figure 1.**
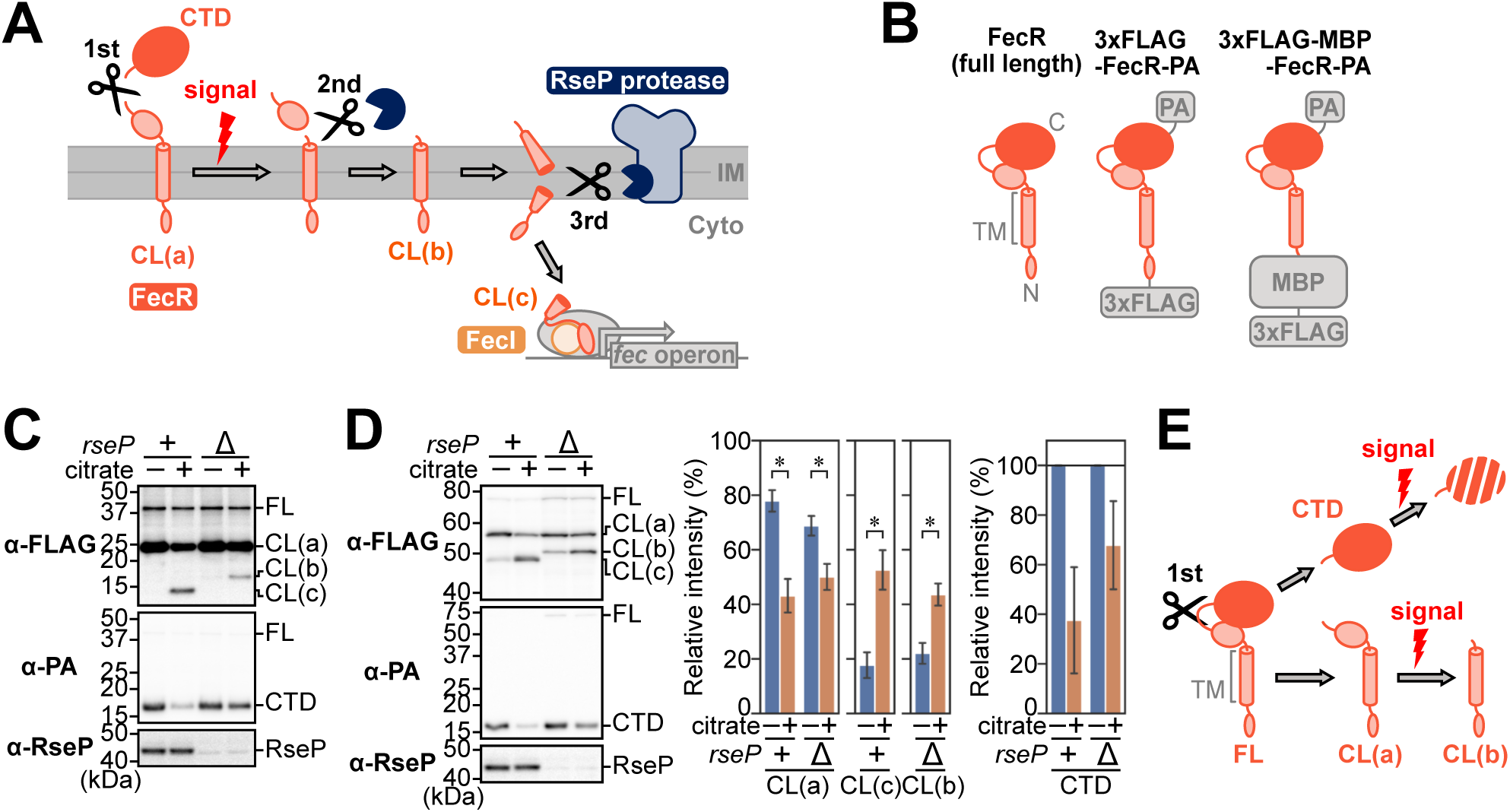
The FecR CTD fragment is degraded depending on the ferric citrate signal. (A) Schematic representation of the FecR processing cascade, adapted with modifications from (23). FecR undergoes an initial autocleavage, generating CL(a) and the CTD fragment (1st cleavage). CL(a) is then truncated at its C-terminal region to produce CL(b) in response to the ferric citrate signal (2nd cleavage). Subsequently, CL(b) is cleaved by the intramembrane protease RseP, yielding CL(c) (3rd cleavage). The released CL(c) interacts with the alternative sigma factor FecI to activate transcription of the *fec* operon. IM and Cyto represent the inner membrane and the cytoplasm, respectively. (B) Schematic representation of the domain organization of FecR and its derivatives used in this study. The transmembrane region (TM) of FecR is shown as a cylinder. (C and D) Cleavage profiles of 3xFLAG-FecR-PA (C) and 3xFLAG-MBP-FecR-PA (D) in response to Na_3_-citrate. *E. coli* strains, YK167 (*rseP*^+^, +) or YK191 (Δ*rseP*, Δ), harboring pYK212 (3xFLAG-FecR-PA) or pYK214 (3xFLAG-MBP-FecR-PA), were grown at 30°C in M9-based medium containing 1 mM isopropyl-*β*-D-thiogalactopyranoside (IPTG) and 10 μM FeCl_3_, with or without 1 mM Na_3_-citrate (citrate) until mid-log phase. Total cellular proteins were acid-precipitated, dissolved in SDS sample buffer containing 10% 2-mercaptoethanol (ME), and analyzed by SDS-PAGE followed by immunoblotting with the indicated antibodies. (C and D, left). FL, CL(a), CL(b), CL(c), and CTD indicate the full-length protein, N-terminal cleavage product CL(a), CL(b), CL(c), and the C-terminal cleavage product CTD fragment, respectively. The band intensities of fragments derived from 3xFLAG-MBP-FecR-PA in indicated strains were quantified (D, middle and right). Band intensities are presented as a percentage of the total intensity of all fragments (D, middle) or normalized to the levels in the citrate-depleted condition (D, right). Data represent the means ± SD from two biologically independent experiments. Student’s *t*-test was carried out to compare the values between the groups. \**P* < 0.05; \*\**P* < 0.01; \*\*\**P* < 0.001; ns, *P* > 0.05. (E) Schematic representation of ferric citrate signal-dependent processing of the FecR CTD and CL(a).

## Results

### The FecR CTD Fragment Generated by the First Cleavage Is Degraded in Response to Ferric Citrate

Our prior findings revealed that FecR cleavage involves a stepwise process, producing a C-terminal domain (CTD) fragment and three distinct N-terminal fragments: CL(a), CL(b), and CL(c) (23) (**Fig. 1A**). This study aimed to investigate the properties and functional role of the CTD fragment produced during the first cleavage of FecR. To detect the CTD fragment, we engineered a derivative of FecR featuring a C-terminal PA tag alongside an N-terminal 3xFLAG tag (3xFLAG-FecR-PA; **Fig. 1B**). This construct retained the ability to transduce the ferric citrate signal (***SI Appendix*, Fig. S1**). The cleavage profile of FecR’s N-terminal region was consistent with previous data supporting our proposed model (**Fig. 1A, C**) (23). Specifically, the level of CL(a) decreased upon citrate addition, indicating that CL(a) undergoes a second cleavage to produce CL(b) in a ferric citrate-dependent manner. CL(b) is further processed by the intramembrane protease RseP to yield CL(c) (**Fig. 1C**). To accurately quantify the cleavage efficiency of degradation-prone, low molecular weight fragments, we designed another derivative where the N-terminal cytoplasmic domain of FecR was replaced with a tightly folded maltose-binding protein (MBP) domain (3xFLAG-MBP-FecR-PA) (**Fig. 1B**). The cleavage patterns of 3xFLAG-MBP-FecR-PA and 3xFLAG-FecR-PA were nearly identical, and the cleavage efficiency could be quantified to support the proposed model (**Fig. 1D**).

Using anti-PA immunoblotting, we detected CTD fragments generated by the first cleavage from both 3xFLAG-FecR-PA and 3xFLAG-MBP-FecR-PA (**Fig. 1C, D**). Interestingly, the levels of these CTD fragments decreased in a citrate-dependent manner (**Fig. 1C, D**), indicating that the CTD fragment undergoes degradation after cleavage in response to ferric citrate (**Fig. 1E**).

### FecR CTD Fragment Is Generated in the Cytoplasm by Autoproteolysis

Next, we analyzed the intracellular behavior of CL(a) and the CTD fragment, focusing on identifying FecR cleavage sites during sequential processing. Mass spectrometry revealed the C-terminal residues of CL(a), CL(b), and CL(c) as Gly-181, Gln-122, and Arg-79/Arg-80, respectively (***SI Appendix*, Fig. S2**). Notably, the first cleavage occurs between Gly-181 and Thr-182, corroborating previous findings obtained through Edman degradation (24). Given that the *Pseudomonas* sigma regulatory factor FoxR undergoes non-enzymatic self-cleavage through an N-O acyl rearrangement (25), we have proposed that FecR similarly self-cleaves (23). However, it remains unclear whether this self-cleavage occurs in the cytoplasm or periplasm after transport.

FecR is inserted into the inner membrane through the Tat pathway, guided by its twin-arginine motif (Arg-79 and Arg-80) (26). To determine whether membrane insertion precedes FecR’s first cleavage, we analyzed fragment production in a strain lacking *tatABC,* which encodes components of the Tat apparatus (27, 28). In this strain, RseP-mediated production of CL(c) was absent, even with citrate supplementation, while ectopic *tatABC* expression restored CL(c) production (**Fig. 2A**). In contrast, CL(a) and the CTD fragment were detected regardless of *tatABC* presence (**Fig. 2A**), indicating that FecR self-cleavage occurs independently of membrane insertion. These results suggest that the first cleavage takes place in the cytoplasm, with the CTD fragment initially localized in this compartment (**Fig. 2B**).

**Figure 2.**
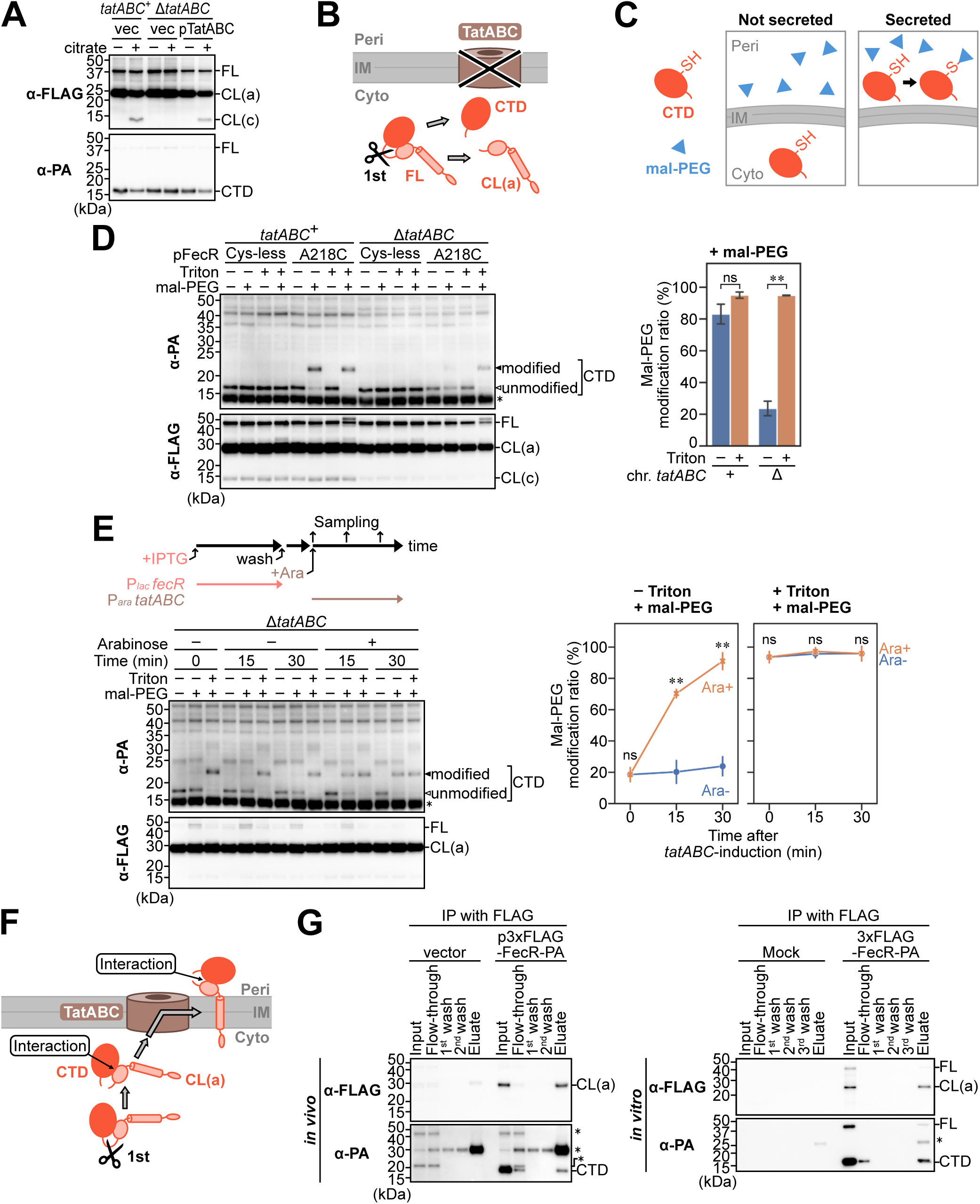
The CTD fragment is exported to the periplasm concurrently with the membrane insertion of CL(a) via the Tat pathway. (A) Cleavage profiles of 3xFLAG-FecR-PA in response to Na_3_-citrate in the presence or absence of TatABC. YK167 (*tatABC*^+^) or YK1196 (Δ*tatABC*) cells harboring pYK212 (3xFLAG-FecR-PA) were transformed with either pSTD689 (vector control, vec) or pYK345 (pTatABC). Cells were grown, and the total cellular proteins were analyzed as described in Fig. 1C. (B) A schematic representation of FecR cleavage in a TatABC-deficient strain. The first cleavage occurs in the cytoplasm, producing CL(a) and the CTD fragment. Peri, IM, and Cyto indicate the periplasm, inner membrane, and cytoplasm, respectively. (C) A schematic representation of the mal-PEG-2k modification assay. The localization of the CTD fragment is assessed by its modifiability with membrane-impermeable mal-PEG-2k. (D) Cellular localization of the CTD fragment assessed by the mal-PEG-2k modification assay in the presence or absence of TatABC. HM1742 (*tatABC*^+^) or YK2238 (Δ*tatABC*) cells harboring pYK1001 (3xFLAG-FecR-PA C271A, pFecR Cys-less) or pYK1025 (3xFLAG-FecR-PA C271A/A218C, pFecR A218C) were grown in Na_3_-citrate-free medium as described in Fig. 1C. Spheroplasts were prepared from these cells and treated with 2 mM mal-PEG-2k (mal-PEG) in the presence or absence of 1% Triton X-100 (Triton). Total proteins were analyzed as described in Fig. 1C. An asterisk indicates a lysozyme band (D, left). The levels of mal-PEG-2k-modification of the CTD fragment were quantified and presented as a percentage of the total levels of the mal-PEG-2k-modified and -unmodified bands. Data represent the means ± SD from two biologically independent experiments. Student’s *t*-test was carried out to compare the values between the groups. \*\**P* < 0.01; ns, *P* > 0.05.(D, right). (E) Tat pathway-dependence of the CTD fragment export assessed by the mal-PEG-2k modification assay. YK2238 (Δ*tatABC*) cells harboring pYK1025, which encodes 3xFLAG-FecR-PA C271A/A218C under the control of the *lac* promoter, were transformed with pYK1000 encoding TatABC under the control of the *araBAD* promoter. A schematic workflow of the experiment is shown (E, upper left). Cells were grown at 30°C in M9-based medium containing 10 μM FeCl_3_ and 0.05% fucose, which inhibits the expression from the *araBAD* promoter, until mid-log phase. FecR expression was induced with 1 mM IPTG for 30 min. Cells were then washed, resuspended in M9-based IPTG-free medium without or with 0.2 % arabinose to induce TatABC expression (indicated as 0 min), and further incubated at 30°C. At the indicated time point, the cells were collected, and the mal-PEG-2k modification assay was performed as described in (D). The immunoblot image (E, lower left) and quantification of mal-PEG-2k-modification (E, right) are shown. The modification level is plotted against the induction time of TatABC expression. Data represent the means ± SD from two biologically independent experiments. Student’s *t*-test was carried out to compare the values between the groups. \*\**P* < 0.01; ns, *P* > 0.05. (F) Schematic representation of the model in which the CTD fragment is exported to the periplasm via the Tat pathway. After the first cleavage, the CTD fragment remains associated with CL(a) and is transported to the periplasm together with the membrane insertion of CL(a). (G) Co-immunoprecipitation assay demonstrating the *in vivo* (G, left) and *in vitro* (G, right) interaction between the CTD fragment and CL(a). Crude membranes were prepared from YK167 cells harboring pTWV228 (vector) or pYK212 (p3xFLAG-FecR-PA), solubilized with Triton X-100, and subjected to immunoprecipitation using anti-FLAG antibody (G, left). For the *in vitro* assay, 3xFLAG-FecR-PA was synthesized using the PURE system in the presence of *n*-Dodecyl-*β*-D-maltoside (DDM) and subjected to immunoprecipitation with anti-FLAG antibody (G, right). Proteins in the input, flow-through, wash, and eluate fractions were analyzed as described in Fig. 1C. Asterisks indicate nonspecific bands.

### FecR CTD Fragment Is Exported to the Periplasm in Association With CL(a) through the Tat System

To determine the subcellular destination of the CTD fragment, we assessed its localization by modifying cysteine residues using the membrane-impermeable reagent methoxypolyethylene glycol 2000 maleimide (mal-PEG; **Fig. 2C**). We introduced a single cysteine at Ala-218 in a Cys-less 3xFLAG-FecR C271A-PA derivative, based on predictions from Alphafold3 (29) that this site is exposed on the surface of the CTD fragment (***SI Appendix*, Fig. S3A**). Both the Cys-less and single-Cys (A218C) mutants retained *fec* signaling function and exhibited cleavage patterns similar to the wild-type (***SI Appendix*, Fig. S3B-D**), without forming disulfide-bonded artifacts (***SI Appendix*, Fig. S3D**). We then quantified the mal-PEG modifiability of this cysteine residue in the spheroplast to assess the membrane translocation ability of the CTD fragment. In the *tatABC*^+^ strain, the CTD fragment from the Cys-less mutant was unmodified, whereas A218C was highly modifiable, even without membrane solubilization by Triton X-100 (**Fig. 2D**). Conversely, in the *tatABC* deletion strain, the CTD fragment was barely modified unless the membrane was solubilized (**Fig. 2D**). The ectopic expression of *tatABC* in the Δ*tatABC* strain resulted in the localization of the CTD fragment to the periplasm (***SI Appendix*, Fig. S3E**). These findings confirm that the CTD fragment is exported to the periplasm through the Tat pathway. As a control, FtsH, a membrane protein with cytoplasmic cysteine residues, was not modified by mal-PEG unless the membrane was solubilized (***SI Appendix*, Fig. S3F**). This result excludes the possibility that overproduction of *tatABC* alters membrane permeability to mal-PEG. To directly demonstrate Tat-dependent transport of the CTD fragment, we developed an experimental system with independent control over FecR and TatABC expression. FecR was transiently expressed in Δ*tatABC* cells under isopropyl-*β*-D-thiogalactopyranoside (IPTG) induction, allowing sufficient time for its self-cleavage in the cytoplasm. Subsequently, *tatABC* expression was induced with arabinose (**Fig. 2E, *upper left***). This analysis revealed that the CTD fragment, once formed in the cytoplasm, is translocated to the periplasm via the Tat pathway (**Fig. 2E, *lower left and right panels***).

Based on these findings, we propose a model in which FecR self-cleavage leaves CL(a) and the CTD fragment associated in the cytoplasm, after which the CTD fragment is exported to the periplasm, accompanied by Tat-mediated membrane insertion of CL(a). To validate this model, we analyzed the interaction between CL(a) and the CTD fragment both in *E. coli* cells (**Fig. 2G, *left panel***) and in a cell-free system (**Fig. 2G, *right panel***). Immunoprecipitation of *E. coli* lysates (**Fig. 2G, *left panel***) and recombinant FecR synthesized using the PURE system, an established cell-free protein synthesis system (30, 31) (**Fig. 2G, *right panel***), demonstrated that these fragments intrinsically interact, independent of cell membranes. These results support a model of protein-protein interaction-mediated co-transport across the inner membrane (**Fig. 2F**).

### FecR CTD Fragment Is Crucial for the Regulation of the Signal-Dependent Cleavage of CL(a) and Activation of FecI

We further explored the role of CTD localization to the periplasm. To this end, we constructed a 3xFLAG-MBP-FecR derivative lacking the CTD (**3xFLAG-MBP-FecR CL(a)-mimic**; **Fig. 3A**) and analyzed its ferric citrate-dependent cleavage. Interestingly, the CL(a)-mimic underwent constitutive cleavage, independent of the ferric citrate signal, yielding CL(b) in Δ*rseP* cells and CL(c) in *rseP*^+^ cells (**Fig. 3B**). Additionally, transcription of the *fec* operon was constitutively induced in the absence of the CTD (**Fig. 3C**), aligning with observations from the isolated expression of the FoxR N-terminal domain (32). These findings indicate that the CTD, when localized to the periplasm, plays a critical role in transmitting the ferric citrate signal to CL(a) and regulating the downstream cleavage cascade necessary for FecI activation (**Fig. 3D**).

**Figure 3.**
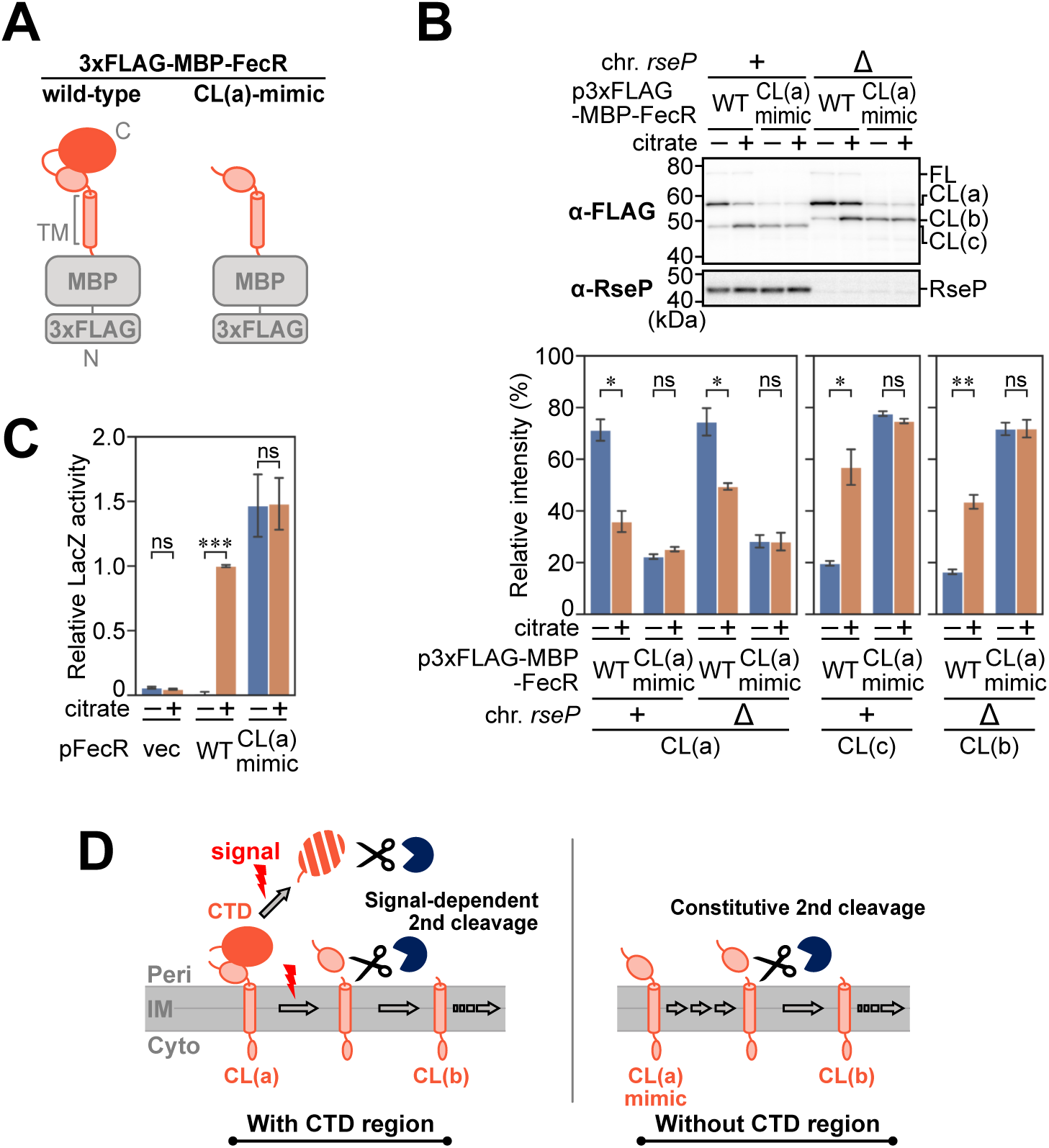
The CTD fragment regulates the signal-dependent cleavage of CL(a). (A) Schematic representation of the domain organization of 3xFLAG-MBP-FecR and its CL(a)-mimic mutant lacking the CTD fragment. (B) Cleavage profiles of the CL(a)-mimic mutant in response to Na_3_-citrate in the presence or absence of RseP. YK167 (*rseP*^+^, +) or YK191 (Δ*rseP*, Δ) cells harboring either pYK147 (p3xFLAG-MBP-FecR WT) or pYK172 (p3xFLAG-MBP-FecR CL(a)-mimic) were grown and total cellular proteins were analyzed. The immunoblot image (B, upper) and the quantified band intensities (B, lower) are presented as described in Fig. 1D. (C) Ability of the FecR CL(a)-mimic mutant to transmit the ferric citrate signal. YK627 (Δ*fecR*) cells harboring pYK149 (P*_fecA_*-*lacZ*) and pSTD343 (*lacI*) were transformed with either pSTD1060 (vector), pYK188 (pFecR WT), or pYK341 (pFecR CL(a)-mimic). Cells were grown at 30°C in M9-based medium containing 1 mM IPTG and 0.1 μM FeCl_3_, with or without 1 mM Na_3_-citrate, until mid-log phase, and their LacZ activities were measured. Relative LacZ activities, normalized to that of the cells harboring pYK188 grown in the same medium with 1 mM Na_3_-citrate, are shown. Data represent the means ± SD from two biologically independent experiments. Student’s *t*-test was carried out to compare the values between the groups. \*\*\**P* < 0.001; ns, *P* > 0.05. (D) Schematic representation of the role of the CTD. The CL(a)-mimic mutant undergoes a constitutive second cleavage, independent of the ferric citrate signal.

### Ferric Citrate Signaling-Mediated Dissociation of the CTD Fragment From CL(a) Triggers the Sequential Cleavage of CL(a)

We next explored the molecular mechanism by which the CTD fragment regulates the second cleavage of FecR. Observations indicated that the CTD fragment physically interacts with CL(a) upon translocation to the periplasm (**Fig. 2**). Based on this, we hypothesized that the CTD fragment inhibits CL(a) cleavage and that its dissociation, triggered by the ferric citrate signal, enables CL(a) cleavage and subsequent degradation of the CTD fragment. To test this hypothesis, we examined whether tethering the CTD fragment to CL(a) through disulfide cross-linking influenced ferric citrate-dependent cleavage. A structural model of the CTD-CL(a) complex, predicted by Alphafold3, suggested that Asn-266 in the CTD and Lys-115 in CL(a) are in close proximity (**Fig. 4A**; ***SI Appendix*, Fig. S4**). We substituted these two residues with cysteine residues, either individually or simultaneously. We reasoned that substituting these residues in FecR would not affect its self-cleavage, as these positions are likely to be distant from the self-cleavage site (**Fig. 4A**). The resulting 3xFLAG-MBP-FecR-PA derivatives were expressed under IPTG induction, followed by disulfide cross-linking induced with the oxidizing agent diamide (**Fig. 4B, *upper panel***). Cells were then washed, incubated in either citrate-containing or citrate-free medium. The total cellular proteins were precipitated, solubilized in the presence or absence of the reducing agent 2-mercaptoethanol (ME), and analyzed by immunoblotting (**Fig. 4B, *middle and lower panels***). Full-length FecR was nearly undetectable in all mutants, confirming that the cysteine substitutions did not impair FecR self-cleavage (**Fig. 4B, *middle panel***). As anticipated, cross-linking between CL(a) and the CTD through disulfide bonds occurred only in mutants with both Lys-115 and Asn-266 cysteine substitutions, strongly inhibiting CL(a) cleavage and CTD degradation (**Fig. 4B, *middle and lower panels***). Exposure of cells to tris(2-carboxyethyl)phosphine (TCEP), a reducing agent, after disulfide cross-linking (**Fig. 4C, *upper panel***) disrupted the disulfide bond, restoring ferric citrate signal-dependent cleavage of CL(a) and degradation of the CTD fragment in a TCEP concentration-dependent manner (**Fig. 4C, *middle and lower panels***). These findings demonstrate that disrupting the interaction between the CTD fragment and CL(a) is essential for CL(a) cleavage and CTD degradation in response to the ferric citrate signal, supporting our hypothesis.

**Figure 4.**
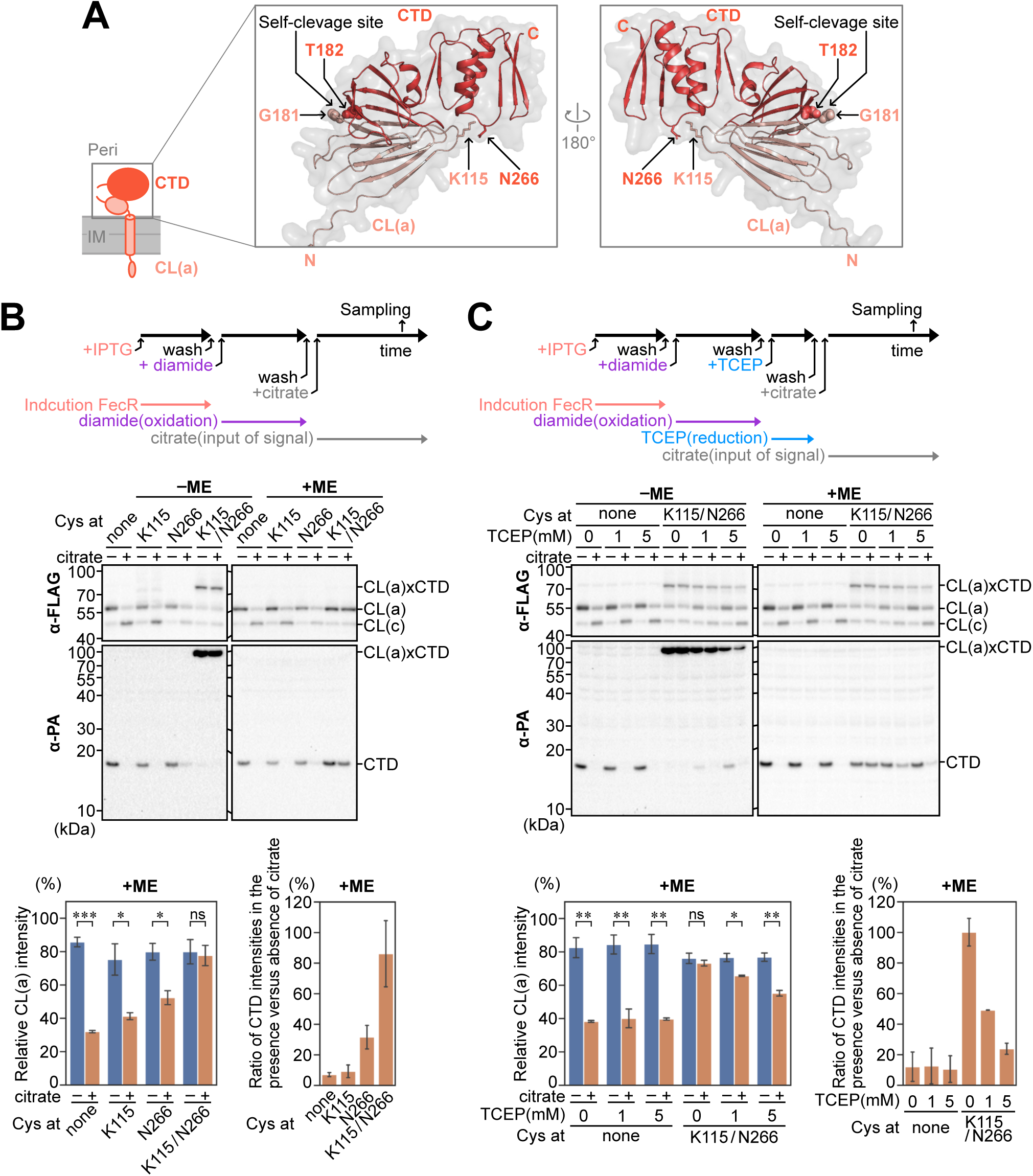
Dissociation of the CTD fragment from CL(a) is essential for ferric citrate signal-dependent CL(a) cleavage and CTD degradation. (A) AlphaFold3-predicted structure of the FecR CL(a)-CTD complex. CL(a) and the CTD fragment are shown in pink and red, respectively. The C-terminal residue of CL(a) (G181) and the N-terminal residue of the CTD fragment (T182) are depicted as sphere models. The side chains of the cysteine-introduced residues (K115 and N266) are presented as stick models. (B) Effect of the disulfide cross-linking between CL(a) and the CTD fragment on CL(a) cleavage and CTD degradation. YK167 cells harboring pYK369 (pFecA) were transformed with plasmids encoding the indicated 3xFLAG-MBP-FecR-PA Cys-less derivatives. FecA is constitutively overproduced from pYK369 to provide sufficient signal transduction to FecR. A schematic workflow of the experiment is shown (B, upper). Cells were grown at 30°C in M9-based medium containing 1 mM IPTG and 10 μM FeCl_3_ until mid-log phase. The cells were then washed and resuspended in IPTG-free M9-based medium containing 10 μM FeCl_3_ and 1 mM diamide. After 1 h of cultivation at 30°C to allow disulfide bond formation, the cells were washed again, resuspended in IPTG-free M9-based medium containing 10 μM FeCl_3_ with or without 1 mM Na_3_-citrate, incubated for additional 1 h at 30°C, and harvested. Total cellular proteins were acid-precipitated, dissolved in SDS sample buffer containing 25 mM N-ethylmaleimide (NEM; to bock free thiol groups), with or without 10% ME, and analyzed by immunoblotting as described in Fig. 1D (B, middle). ‘CL(a)xCTD’ indicates the disulfide cross-linked product. Band intensities of CL(a) and CL(c) (B, lower left) and the CTD fragment (B, lower right) under reducing conditions (+ME) were quantified as described in Fig. 1D. (C) Effect of reducing the disulfide bond between CL(a) and the CTD fragment on CL(a) cleavage and CTD degradation. Cells were grown and treated as described in (B). After treatment with diamide, the cells were washed, resuspended in IPTG-free M9-based medium containing 10 μM FeCl_3_ and the indicated concentration of tris(2-carboxyethyl)phosphine (TCEP), and incubated at 30°C for 30 min. The cells were then washed again, resuspended in IPTG-free M9-based medium containing 10 μM FeCl_3_ with or without 1 mM Na_3_-citrate, incubated for 1 h at 30°C, and harvested. Proteins were analyzed as described in (B), and the results are shown in the same format.

### TonB-Dependent Transporter FecA Is Involved in FecR CTD Dissociation From CL(a), Facilitating the Proteolysis-Mediated Ferric Citrate Signaling Pathway

Fec signaling is initiated by the binding of ferric citrate to the outer membrane transporter FecA (11, 33). Iron transport across the outer membrane relies on the interaction between FecA and TonB in the periplasm. TonB, in turn, is associated with the ExbBD motor complex in the inner membrane (11, 34). As a member of the TBDT family and a homolog of the TBDT FhuA (35), FecA plays a critical role in this process. Based on prior evidence that FecR and FecA interact in signal transduction (15, 16), we hypothesized that mechanical force generated by the TonB-ExbBD complex is transmitted to FecA and subsequently relayed to the FecR CTD, leading to the dissociation of the CTD fragment from CL(a) .

Using AlphaFold3, we predicted a complex structure comprising FecR CL(a), FecR CTD, FecA, and TonB, suggesting that these components can interact simultaneously (**Fig. 5A**; ***SI Appendix*, Fig. S5A**). The model highlights the *β*-augmentation interaction between TonB and the TonB box of FecA, a conserved periplasmic domain in TBDTs with high affinity for TonB’s C-terminal domain (20, 21) (**Fig. 5A, *upper right panel***). Notably, the signal-dependent production of FecR CL(c) occurred only in the presence of TonB (**Fig. 5B**), indicating that TonB is essential for progressing the FecR cleavage cascade but not required for the initial cleavage producing CL(a) and the CTD fragment. To further investigate the role of TonB-dependent mechanical force in the FecR cleavage cascade, we conducted experiments using carbonyl cyanide *m*-chlorophenyl hydrazone (CCCP), a protonophore that disrupts the PMF (**Fig. 5C**). FecR expression was first induced with IPTG, after which the cells were washed, treated with CCCP, washed again, and incubated with citrate in the continued presence of CCCP. We observed that citrate-induced cleavage of CL(a) was almost entirely inhibited by CCCP, demonstrating that PMF-driven TonB mechanical motion is indispensable for the second cleavage of FecR. The AlphaFold3 model also predicts that the FecR CTD fragment interacts with the N-terminal region of FecA through a *β*-augmentation mechanism (**Fig. 5A, *right middle and lower panels, SI Appendix*, Fig. S5A**). To examine the functional importance of this interaction, we deleted the N-terminal regions of FecA, including the *β*-strand region (“Interaction *β*-strand”) predicted to serve as the interaction interface with the FecR CTD (***SI Appendix*, Fig. S5A**). These deletions prevented the ferric citrate signal-dependent cleavage of FecR CL(a) **(*SI Appendix*, Fig. S5B)**. Furthermore, replacement of residues in this *β*-strand with proline almost completely abolished ferric citrate-dependent CL(a) cleavage and CTD degradation (**Fig. 5D**). These results highlight the critical role of FecA-FecR CTD *β*-augmentation in the FecR cleavage cascade. Based on these findings, we propose a model where FecA, energized by the TonB-ExbBD motor complex, triggers CTD dissociation from CL(a) and CTD degradation, initiating sequential cleavage of FecR CL(a) and cytoplasmic signal transduction.

**Figure 5.**
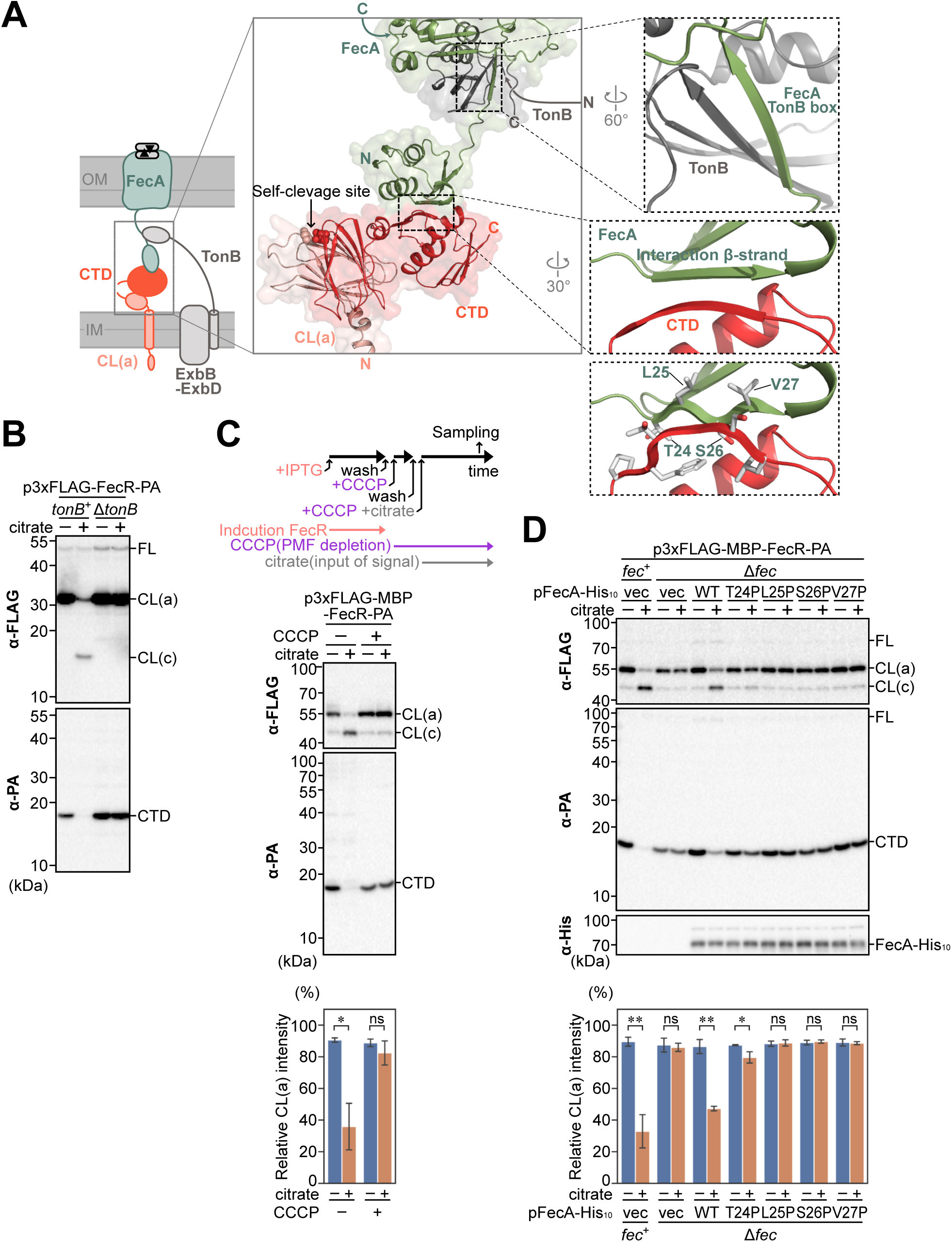
The FecA transporter, powered by the TonB motor, mediates the dissociation of the CTD fragment from CL(a), enabling ferric citrate signal-dependent CL(a) cleavage. (A) AlphaFold3-predicted structure of the CL(a)-CTD-FecA-TonB supercomplex. FecR CL(a), the CTD fragment, FecA, and TonB are shown in pink, red, green, and gray, respectively. Enlarged views of the interaction sites between TonB and FecA (upper dashed box) and between FecA and the FecR CTD fragment (middle and lower dashed boxes) are shown. (B) Cleavage profiles of the FecR protein in the presence or absence of TonB. YK167 (*tonB*^+^) or YK2234 (Δ*tonB*) cells were transformed with pYK212 (p3xFLAG-FecR-PA). Cells were grown, and proteins were analyzed as described in Fig. 1C. (C) Effect of the depletion of PMF on CL(a) cleavage and CTD degradation. YK167 cells harboring pYK367 (pFecA) were transformed with pYK214 (p3xFLAG-MBP-FecR-PA). A schematic workflow of the experiment is shown (C, upper). Cells were grown at 30°C in M9-based medium containing 1 mM IPTG and 10 μM FeCl_3_ until mid-log phase. The cells were then washed and resuspended in IPTG-free M9-based medium containing 10 μM FeCl_3_ and 100 μM carbonyl cyanide *m*-chlorophenyl hydrazone (CCCP). After 15 min of incubation at 30°C, the cells were washed again, resuspended in M9-based IPTG-free medium containing 10 μM FeCl_3_ and 100 μM CCCP with or without 1 mM Na_3_-citrate, incubated for additional 1 h at 30°C, and the total cellular proteins were analyzed. The immunoblot image (C, middle) and the quantified band intensities (C, lower) are presented as described in Fig. 1D. (D) Effect of FecA mutations on the predicted FecA-FecR CTD interaction, assessed by FecR cleavage. YK167 (*fecA*^+^) or YK1602 (Δ*fecA*) cells harboring pYK214 (p3xFLAG-MBP-FecR-PA) were transformed with plasmids encoding the indicated FecA-His_10_ derivatives. Cells were grown, and total cellular proteins were analyzed as described in Fig. 1D.

## Discussion

Early studies identified TonB as essential for Fec signaling (11, 34), yet the molecular mechanism remained elusive. Here, we propose a comprehensive model for the Fec signaling pathway **(Fig. 6)**. FecR undergoes an initial self-cleavage in the cytoplasm after translation, generating CL(a) and the CTD fragment, which remain associated. During CL(a) membrane insertion via the Tat system, the CTD fragment is co-transported to the periplasm. Ferric citrate binding to FecA exposes its TonB box to the periplasm, enabling interaction with the TonB C-terminal domain of TonB, as demonstrated by many earlier studies [reviewed by (20, 21)]. This interaction induces a conformational change in FecA, promoting its N-terminal signaling domain (NTSD) interaction with the FecR CTD. PMF-driven mechanical force exerted by TonB dissociates the FecR CTD from CL(a), facilitating its degradation and enabling the second cleavage of CL(a) into CL(b). Subsequently, RseP cleaves CL(b), generating CL(c), which translocates to the cytoplasm and activates fec operon transcription via the FecI sigma factor.

**Figure 6.**
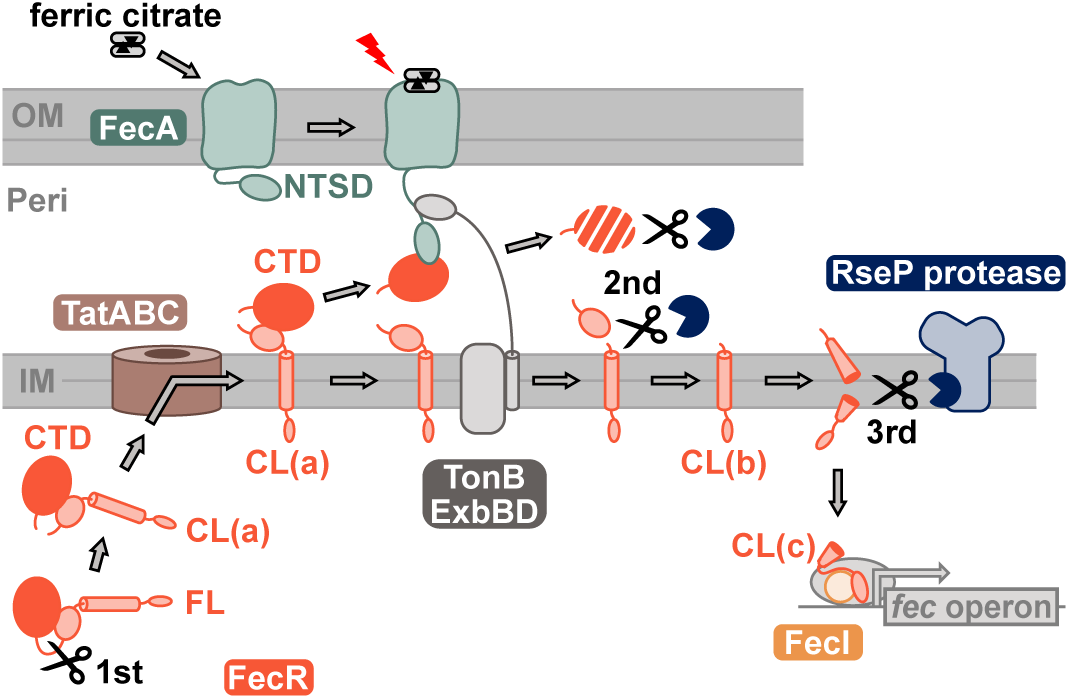
Model of the Fec signaling mechanism. The TonB-dependent transporter FecA mediates the progression of the FecR cleavage cascade in response to the ferric citrate signal. See the text for a detailed description.

FecA TBDT has dual roles: (i) transporting iron carriers and (ii) recognizing ferric citrate to initiate signal transduction. While the TonB-dependent gating mechanism of outer membrane transporters has been extensively studied, leading to a model in which TonB-mediated unfolding the plug domain of TBDTs is powered by the ExbBD motor (20, 21), TonB’s role in TBDT-mediated signal transduction remains poorly understood. We propose that TonB-powered dissociation of the FecR CTD from CL(a), mediated by FecA through its TonB box, is the critical event triggering the sequential cleavages of CL(a). These cleavages ultimately generate the sigma-interacting fragment CL(c). Thus, the TonB-FecA interaction establishes a vital link between substrate transport and signal transduction.

In our model, the dissociation of the FecR CTD from CL(a) is central, though we cannot rule out the possibility that structural changes at the fragment interface facilitate CL(a) cleavage. Interestingly, we observed that FecR CTD undergoes degradation by as-yet-unknown periplasmic protease(s) in response to the ferric citrate signal, a process reminiscent of the degradation of FoxR in response to ferrioxamine (32). This degradation likely occurs when structural changes in FecR CTD, induced by its release from CL(a), trigger protease recognition. Degrading CTD may prevent its re-association with CL(a), ensuring the unidirectional progression of the signaling cascade. Alternatively, if FecR CTD remains associated with FecA post-signal reception, its degradation might allow FecA to reset for subsequent signaling cycles. Identifying the periplasmic protease(s) responsible for producing CL(b) is critical to elucidating the cleavage cascade. Despite comprehensive screening by disrupting candidate genes (***SI Appendix*, Fig. S6**), no single gene was found essential for CL(a) cleavage. This could mean that multiple enzymes with overlapping functions are involved, necessitating further investigation.

A similar cleavage-dependent signal transduction mechanism has been reported in the σ^Ε^-mediated envelope stress response in *E. coli*. Here, DegS-catalyzed cleavage of RseA, an anti-sigma factor, initiates a cascade completed by RseP-mediated cleavage, releasing σ^E^ into the cytoplasm (36). The periplasmic protein RseB plays a regulatory role by inhibiting DegS-mediated proteolysis through its binding to the periplasmic domain of RseA (37, 38). RseB inhibition is relieved when RseB dissociates from RseA in response to dysfunction in outer-membrane biogenesis (39), enabling the RseA cleavage cascade to proceed, consequently activating the transcription of stress-response genes. However, RseB alone plays a limited role in the regulation in this scheme, whereas FecR CTD plays a very critical role in Fec signaling (40).

The Tat system is a well-characterized transporter that exports folded proteins across the cytoplasmic membrane into the periplasm (27). This system often ensures quality control for substrate proteins, particularly those with redox cofactors (28, 41). Some redox enzyme complexes co-transport partner proteins lacking a twin-arginine motif through association with Tat signal-holding proteins (28). For instance, nickel-containing hydrogenases, which include a subunit devoid of an export signal, are transported through the Tat system by the hitchhiker mechanism (42). In this study, we experimentally validated the Tat-mediated transport of two peptides that fold and associate in the cytoplasm. This is the first to demonstrate that two peptides, generated through self-cleavage of a single protein, can function as Tat substrates. Furthermore, to the best of our knowledge, this is the first observation of the co-translocation of multiple proteins via the Tat system outside the context of redox enzymes. The two fragments of FecR, CTD and CL(a), are associated with each other as folded conformations prior to membrane translocation and maintain this associated form in the periplasm. These characteristics would be key requirements for their tight regulation and prompt response to ferric citrate, which triggers successive cleavage. Without mutual association, both fragments would exhibit intrinsic instability. We speculate that the self-cleavage of FecR into these fragments, rather than encoding them at separate loci, ensures proper folding and function as a complex.

Iron is a critical element for nearly all life forms, yet its low solubility under physiological conditions poses significant challenges for cellular acquisition (43). Many pathogenic bacteria have coevolved with eukaryotic hosts to adapt to iron-limited environments. Intracellular iron in host cells is primarily stored in ferritin or bound to metalloproteins, while extracellular iron is largely sequestered in heme, complexed with hemoglobin (44, 45). To overcome these limitations, gram-negative bacteria employ conserved iron uptake systems (33, 46), which are closely linked to their virulence (47–53). Understanding the molecular mechanisms of signal transduction in these systems is essential for elucidating bacterial pathogenicity and environmental adaptation. Building on our recent finding of the sequential cleavages of the sigma regulator FecR (23), we have unraveled the core mechanism of Fec signaling, revealing the unique intracellular behaviors of this protein in association with the TonB-dependent transporter FecA.

## Materials and Methods

### Bacterial Strains, Plasmids, Media, and Antibodies

The *E. coli* K12 strains and plasmids used in this study are listed in ***SI Appendix, Table S1***. Detailed descriptions of the construction of the strains and plasmids, as well as the media and antibodies used in this research, are provided in ***SI Appendix, Materials and Methods***.

### SDS-PAGE Assay and Immunoblotting Analysis

SDS-PAGE and immunoblotting were performed essentially as described previously (23, 54). Further details are provided in ***SI Appendix, Materials and Methods***.

### Mal-PEG Modification Assay

Spheroplasts were treated with or without 2 mM mal-PEG-2k and/or 1% Triton X-100 for 30 minutes at 4°C. Unreacted mal-PEG-2k was quenched by the addition of 100 mM dithiothreitol (DTT), and proteins were precipitated using trichloroacetic acid. The resulting samples were analyzed through SDS-PAGE and immunoblotting assays. Additional procedural details are provided in ***SI Appendix, Materials and Methods***.

### Immunoprecipitation Assay

Total membrane extracts from cells expressing 3xFLAG-FecR-PA, or 3xFLAG-FecR-PA synthesized using the PURE*frex*^®^ 1.0 cell-free protein synthesis kit (Gene Frontier) (54), were subjected to immunoprecipitation with anti-FLAG M2 affinity gel (Merck). Further methodological details are provided in ***SI Appendix, Materials and Methods***.

### Structural Prediction With AlphaFold3

Structural models were generated using AlphaFold3 (29). Further details are provided in ***SI Appendix, Materials and Methods***.

### *β*-Galactosidase (LacZ) Activity Assay

LacZ activity in cells harboring the reporter plasmid pYK149 (P*_fecA_*-*lacZ*) was measured as previously described (23). Further details are available in ***SI Appendix, Materials and Methods***.

## Acknowledgments

We sincerely thank Hiroyuki Mori for his stimulating discussions and valuable advice. We also thank Kazue Kanehara for constructing the strain KK379, Hanayo Hirose and Mariko Yoshioka for their technical assistance, Kayo Abe, Satomi Koshiba, and Michiyo Sano for their secretarial support, the other laboratory members for their valuable discussion, and National BioResource Project (NBRP)-*E. coli* at National Institute of Genetics, Japan for providing *E. coli* strains. This research was funded by grants from the Japan Society for the Promotion of Science (JSPS) KAKENHI to T.Y. (JP23KJ1581, JP24H01370, and JP24K17817), R.M. (JP22K15061), T.T. (JP21H05155), and Y.A. (JP22H02571/23K23835), and Institute for Fermentation, Osaka (IFO) to T.Y. (Y-2023-2-023).

## Data availability statement

All study data are included in the article and/or supporting information.

