## Supplementary text and figures for "Cleavage cascade of the sigma regulator FecR orchestrates TonB-dependent signal transduction"

**This PDF file includes:**

Supporting text
Figures S1 to S6
Tables S1
Legends for Datasets S1 to S3
SI References

**Other supporting materials for this manuscript include the following:**

Datasets S1 to S3

### Supplementary Materials and Methods

#### Media and bacterial culture.

M9 synthetic medium (without $CaCl_2$) supplemented with 2 µg/mL thiamine, 20 kinds of amino acids (20 µg/mL each), and either 0.4% glucose or 0.2% maltose with 0.4% glycerol were used for *E. coli* cultivation. To induce *lac* promoter/operator-controlled gene expression in cells grown in M9-based medium containing glucose, 1 mM cAMP was added to the medium in addition to 1 mM IPTG. Antibiotics, including ampicillin (50 µg/ml), chloramphenicol (20 µg/ml), and/or spectinomycin (50 µg/ml) were used for selecting transformants and for growing plasmid-harboring cells. Bacterial growth was monitored using either a Miniphoto 518R (660 nm; TAITEC) or a Miniphoto 620 (620 nm; TAITEC).

#### Antibodies.

Mouse monoclonal anti-FLAG M2 antibody (Merck) and anti-His tag antibody (MBL), rat monoclonal anti-PA antibody (Fujifilm), and rabbit polyclonal anti-FtsH antibody (1) and anti-RseP antibody (2) were used for immunoblotting. For the immunoprecipitation assay, anti-FLAG M2 affinity gel (Merck) was used. The following secondary antibodies were used for immunoblotting; Goat Anti-Rabbit IgG (H + L)-HRP conjugate (Bio-Rad), Goat anti-Rabbit IgG (H+L) secondary antibody, HRP (Thermo Fisher), Goat Anti-Mouse IgG (H + L)-HRP conjugate (Bio-Rad), Goat anti-Mouse IgG (H+L) secondary antibody, HRP (Thermo Fisher), Mouse TrueBlot® ULTRA: Anti-Mouse Ig HRP (Rockland) or Anti-Rat IgG (whole molecule)−Peroxidase antibody produced in rabbit (Merck).

#### SDS-PAGE and immunoblotting analysis.

Proteins were dissolved in SDS sample buffer, separated by SDS-PAGE, and electrotransferred onto Immobilon-P membrane (Merck). After blocking, the membrane was incubated with an appropriate primary antibody. For anti-RseP immunoblotting, anti-RseP antibodies were preincubated with whole-cell lysates of AD1840 (the Δ*rseA* Δ*rseP* Δ*degS* strain) at 4 °C for 1 h to reduce background noise, as described previously (3, 4). The membrane was then washed and incubated with an HRP-conjugated secondary antibody. After further washing, proteins were visualized using detection reagents (ECL Western Blotting Detection Reagents (Cytiva), ECL Prime Western Blotting Detection Reagents (Cytiva), Chemi-Lumi One (Nacalai Tesque), or Western Lightning Plus ECL (Revvity)) and chemiluminescence image analyzers (LAS3000 mini lumino-image analyzer (Cytiva), LAS4000 mini lumino-image analyzer (Cytiva), FUSION Solo S (VILBER), or ChemiDoc XRS+ (Bio-Rad)).

#### Mal-PEG modification assay.

Spheroplasts were prepared as follows; cells were washed, resuspended in spheroplast buffer (30 mM Tris-HCl (pH 8.1), 20% sucrose) supplemented with 100 µg/mL lysozyme and 10 mM EDTA, and incubated for 1 h on ice. After the addition of 20 mM MgCl, spheroplasts were treated with mal-PEG-2k and analyzed as described in the Materials and Methods section in the main text.

#### Immunoprecipitation assay.

The co-immunoprecipitation assay for the *in vivo* interaction between FecR fragments was performed as follows: cells expressing 3xFLAG-FecR-PA were harvested, suspended in 10 mM Tris-HCl (pH 8.1) supplemented with 1 mM EDTA, 1 mM Pefablock (Merck), and Protease Inhibitor Cocktail (Nacalai Tesque), and disrupted by sonication. Total membranes were collected by ultracentrifugation (110,000 g for 60 min at 4 °C) and resuspended in buffer A (50 mM Tris-HCl (pH 8.1), 150 mM NaCl, 10% glycerol). To solubilize the membrane, fivefold buffer B (50 mM Tris-HCl (pH 8.1), 150 mM NaCl, 10% glycerol, 1% Triton X-100, 1 mM EDTA, 1 mM Pefablock (Merck), Protease Inhibitor Cocktail (Nacalai Tesque)) were added to the suspension, followed by incubation on ice for 1 h. After clarification by centrifugation, the supernatant was incubated with anti-FLAG M2 affinity gel (Merck) overnight at 4 °C with rotation. The immunocomplexes were collected, washed with buffer B, suspended in 2-mercaptoethanol (ME)-free SDS sample buffer, and boiled. After centrifugation, the supernatant was supplemented with 10% ME and subjected to SDS-PAGE and immunoblotting analyses.

The synthesis of 3xFLAG-FecR-PA using the PURE system was performed as follows: a DNA fragment encoding 3xFLAG-FecR-PA was amplified by two-step PCR reactions. The first PCR was performed using pYK212 as a template and the primer pair P101/P102. The second PCR was carried out using the product of the first PCR as a template and the primer pair P102/P103. The resulting DNA template was used in the PURE*frex*® 1.0 reaction in the presence of 0.01 % *n*-dodecyl-*β*-D-maltoside (DDM) and 1/80 volume of recombinant RNase inhibitor (Takara Bio), and the reaction was incubated at 37 °C for 3 h. The reaction was terminated by the addition of 0.125 mg/mL chloramphenicol. After clarification by centrifugation, the supernatant containing the synthesized FecR fragments was diluted 70-fold with buffer C (50 mM Tris-HCl (pH 7.4), 150 mM NaCl, 1 % Triton X-100). The interaction between *in vitro* synthesized FecR fragments was analyzed using anti-FLAG M2 affinity gel (Merck) in a manner similar to the *in vivo* analysis. The immunocomplexes were collected, washed with buffer C and then with buffer D (50 mM Tris-HCl (pH 7.4), 150 mM NaCl), suspended in ME-free SDS sample buffer, and boiled. After centrifugation, the supernatant was supplemented with 10% ME and subjected to SDS-PAGE and immunoblotting analyses.

**Structure prediction using AlphaFold3.**
The following protein sequences were submitted to the AlphaFold3 program via the AlphaFold server (https://alphafoldserver.com/). The best-scoring model was used for analysis and is presented in a cartoon representation using the PyMOL Molecular Graphics System, Version 3.0 (Schrödinger, LLC). The structural model of the FecR CL(a)-CTD fragment complex shown in Fig. 4A and *SI Appendix*, Fig. S4 was generated using the protein sequences of FecR CL(a) (Uniprot ID: P23485, residues R80 to G181), and the FecR CTD fragment (Uniprot ID: P23485, residues T181 to L317). The resulting model is provided in Dataset S1. The structural model of the FecR CL(a)-CTD fragment-FecA-TonB supercomplex shown in Fig. 5A and *SI Appendix*, Fig. S5A was generated using the protein sequences of FecR CL(a) (Uniprot ID: P23485, residues R80 to G181), FecR CTD fragment (Uniprot ID: P23485, residues T181 to L317), TonB (Uniprot ID: P02929, residues A155 to Q239), and FecA (Uniprot ID: P13036, residues A1 to D222 of its mature form). The resulting model is provided in Dataset S2. The structural model of the full-length form of FecR shown in Fig. S3A was generated using the protein sequence of FecR (Uniprot ID: P23485, residues M1 to L317). The resulting model is provided in Dataset S3.

***β*-galactosidase (LacZ) activity assay.**
LacZ activity of cells harboring the reporter plasmid pYK149 ($P_{fecA}$-*lacZ*) was measured as described previously (3). Briefly, cultures were mixed with Reporter 5xLysis buffer (Promega), frozen at −80 °C for at least 1 h, and thawed by incubation at 37 °C for 30 min in a clear 96-well plate. An equal volume of Z-buffer (60 mM $Na_2HPO_4 \cdot 7H_2O$, 40 mM $NaH_2PO_4 \cdot H_2O$, 10 mM KCl, 1 mM $MgSO_4 \cdot 7H_2O$, 40 mM ME) supplemented with 1.32 mg/ml 2-nitrophenyl *β*-D-galactopyranoside (ONPG) was added to the lysate and incubated at room temperature. Relative LacZ activity was calculated as follows: for measurements in *SI Appendix*, Fig. S1A, absorbance at 420 nm and 550 nm was recorded over time using the Viento Nano microplate reader (BioTek Instruments). “Raw LacZ activity” was calculated as: ($A_{420}$-1.75 x $A_{550}$)/(incubation time (min)). For the other figures, absorbance at 405 nm was recorded over time using the Multiskan FC (Thermo Fisher), and “raw LacZ activity” was calculated as: ($A_{405}$)/(incubation time (min)). “Raw LacZ activity” was normalized by dividing each value by that of a standard cell sample (CU141 cells cultured in M9-based medium) and by the $A_{660}$ of the bacterial culture at the time of collection, giving the “corrected LacZ activity”. Relative LacZ activity was calculated by dividing the “corrected LacZ activity” by that of the corresponding control, as described in the figure legends.

**Mass spectrometric analysis.**
Total cellular proteins of *rseP*$^+$ or Δ*rseP* cells expressing $His_{10}$-MBP-FecR were acid-precipitated and solubilized with buffer E (50 mM Tris-HCl (pH 8.1), 100 mM NaCl, 1 mM phenylmethanesulfonyl fluoride (PMSF)) containing 1 % SDS. After clarification by centrifugation, the supernatant was diluted 10-fold with buffer E, and incubated with TALON metal affinity resin (Takara Bio) for 3 h with rotation. The resin was collected, washed with buffer E containing 0.1 % SDS, suspended in SDS

sample buffer, and shaken vigorously for 10 min at room temperature. After centrifugation, the supernatant was subjected to SDS-PAGE and analyzed by silver staining.

The bands separated by SDS-PAGE were excised, reduced with dithiothreitol (DTT), and alkylated with acrylamide. The treated gel slices were then digested with either trypsin (TPCK-treated, Worthington Biochemical Co.) or endoproteinase Asp-N (Roche). The resulting peptides were analyzed by LC-MS/MS using an Easy-nLC 1200 system coupled with a Q Exactive HF-X hybrid quadrupole-Orbitrap mass spectrometer (Thermo Fisher Scientific).

Peptide separation was performed on a reversed-phase nano-spray column (NTCC-360/75-3-105, NIKKYO Technos) with solvent A (0.1% formic acid) and solvent B (80% acetonitrile with 0.1% formic acid). A linear gradient increasing solvent B from 0% to 40% over 10 minutes was applied. MS and MS/MS data were acquired using the Top10 data-dependent acquisition method. The acquired data were processed using Proteome Discoverer 3.0 (Thermo Fisher Scientific) and Sequest HT 1.17 (Thermo Fisher Scientific). Database searches were performed against an in-house database using the following parameters: enzyme = trypsin (semi-specific) or Asp-N (semi-specific); maximum missed cleavage sites = 2; minimum peptide length = 6; maximum peptide length = 150; dynamic modifications = oxidation (M), propionamide (C); peptide mass tolerance = ±15 ppm; fragment mass tolerance = ±0.03 Da.

Peptides containing cleavage sites were verified by extracting MS chromatograms in Qual Browser (Thermo Fisher Scientific) based on the presumed cleavage sites determined from SDS-PAGE band mobility and comparing these with peptides identified by Sequest HT. Peptides not identified by Sequest HT were manually annotated using Qual Browser.

**Construction of strains.**
YK1131 (Δ*tatABC*::*kan*) and YK1129 (Δfec*IRABCDE*::*kan*) were constructed by deleting the *tatA*-*tatB*-*tatC* or *fecI*-*fecR*-*fecA*-*fecB*-*fecC*-*fecD*-*fecE* of BW25113 by the one-step method described by Datsenko and Wanner (2000) (5) using pKD13, pKD46, and the primer pairs P1/P2 and P3/P4, respectively. YK2234 and YK2238 were constructed by transferring the Δ*tonB*::*kan* and Δ*tatABC*::*kan* markers from JW5195 and YK1131, respectively, into YK167 and HM1742 by P1 transduction, respectively. YK1196 and YK1602 were constructed by transferring the Δ*tatABC*::*kan* and Δ*fecIRABCDE*::*kan* markers from YK1131 and YK1129, respectively, into YK167 by P1 transduction, followed by the deletion of the *kan* cassette using pCP20 (5). KK379 (Δ*rseA*::*cat*) was a derivative of CU141 and was constructed by P1 transduction as described previously (6).

**Construction of plasmids.**
pYK336 [pTWV228 3xFLAG-FecR CL(a)-mimic (T182stop)] was constructed by replacing the codon for the T182 of FecR on pYK130 with the ochre codon by site-directed mutagenesis. pYK341 [pSTD1060 3xFLAG-FecR CL(a)-mimic (T182stop)] was constructed by cloning the EcoRI/HindIII fragment of pYK336 into the same site of pSTD1060. pYK212 (pTWV228 3xFLAG-FecR-PA) and pYK214 (pTWV228 3xFLAG-MBP-FecR-PA) were constructed as follows. The PA tag-encoding insert fragment was prepared by annealing oligonucleotides P11/P12. Vector fragments were PCR-amplified from pYK130 or pYK147 using the primer pair P13/P14 and then ligated with the insert fragment using the In-Fusion® HD cloning kit (Clontech) to generate pYK212 and pYK214, respectively. pYK1001 [pTWV228 3xFLAG-FecR (C271A)-PA], pYK1025 [3xFLAG-FecR (C271A, A218C)-PA], pYK1048 [pTWV228 3xFLAG-FecR (C271A, K115C)-PA], pYK1060 [pTWV228 3xFLAG-FecR (C271A, N266C)-PA], and pYK1079 [pTWV228 3xFLAG-FecR (C271A, K115C, N266C)-PA] were constructed from pYK212 by site-directed mutagenesis using a pair of appropriate primers. pYK238 (pSTD1060 3xFLAG-FecR-PA), pYK1006 [pSTD1060 3xFLAG-FecR (C271A)-PA], and pYK1102 [pSTD1060 3xFLAG-FecR (C271A, A218C)-PA] were constructed by cloning the EcoRI/HindIII fragment of pYK212, pYK1001, and pYK1025, respectively, into the same site of pSTD1060. pYK2004 [pTWV228 3xFLAG-MBP-FecR (C271A)-PA], pYK2006 [pTWV228 3xFLAG-MBP-FecR (C271A, K115C)-PA], pYK2008 [pTWV228 3xFLAG-MBP-FecR (C271A, N266C)-PA], and pYK2010 [pTWV228 3xFLAG-MBP-FecR (C271A, K115C, N266C)-PA] were constructed as follows. Insert fragments encoding the corresponding derivatives of FecR were PCR-amplified from pYK1001, pYK1048, pYK1060, or pYK1079 with the primer pair P15/P16, and ligated into the BamHI/SalI site of pYK214 using the In-Fusion® HD cloning kit (Clontech) to generate pYK2004, pYK2006, pYK2008 and pYK2010, respectively. pYK202 (pTWV228 His$_{10}$-

MBP-FecR) was constructed by cloning annealed oligonucleotides P17/P18 into the SacI/KpnI site of pYK147. pYK364 (pTWV228 FecA) was constructed as follows. A fragment containing the complete ORF of *fecA* with an EcoRI site plus an improved Shine-Dalgarno sequence at the 5' end and a HindIII site at the 3' end was amplified from the MC4100 chromosome by colony PCR using the primer pair P19/P20. After digestion with EcoRI and HindIII, the fragment was cloned into the same site of pTWV228. The obtained plasmid has an unexpected F709L mutation, which was corrected by site-directed mutagenesis to generate pYK364. pYK367 (pSTD689 FecA) and pYK369 (pSTD689c FecA) were constructed by cloning the EcoRI/HindIII fragment of pYK364 into the same site of pSTD689 and pSTD689c, respectively. pYK2023 (pSTD689 FecA-His$_{10}$) was constructed using the In-Fusion® HD cloning kit (Clontech) by self-ligating the fragment that was PCR-amplified from pYK367 using the primer pair P21/P22. pYK2026, pYK2027, pYK2028, and pYK2029 were constructed from pYK2023 by site-directed mutagenesis using a pair of appropriate primers. pYK2030, pYK2031, and pYK2032 were constructed using the In-Fusion® HD cloning kit (Clontech) by self-ligating the fragment that was PCR-amplified from pYK2023 using primer pairs P23/P24, P25/P26, and P27/P28, respectively. pYK301 (pTWV228 TatA, TatB, TatC) was constructed as follows. A fragment containing the complete ORF of *tatABC* with a KpnI site plus the Shine-Dalgarno sequence of *tatA* at the 5' end and a SalI site at the 3' end was amplified from the MC4100 chromosome by colony PCR using the primer pair P29/P30. After digestion with KpnI and SalI, the fragment was cloned into the same site of pTWV228. pYK345 (pSTD689 TatA, TatB, TatC) and pYK1000 (pBAD33 TatA, TatB, TatC) were constructed by cloning the KpnI/SalI fragment of pYK301 into the same site of pSTD689 and pBAD33, respectively. pYK1100 (pSTD689 TonB) was constructed as follows. An insert fragment encoding the complete ORF of *tonB* were PCR-amplified from the MC4100 chromosome by colony PCR using the primer pair P31/P32. A vector fragment was PCR-amplified from pYK367 using the primer pair P33/P34, and then ligated with the insert fragment using the In-Fusion® HD cloning kit (Clontech) to generate pYK1100.

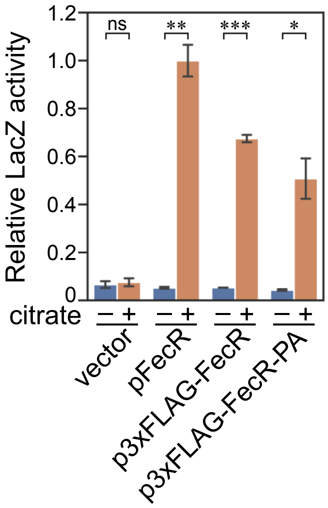


**Fig. S1. Ability of FecR derivatives to transmit the ferric citrate signal.**

YK627 (Δ*fecR*) cells harboring pYK149 ($P_{fecA}$-*lacZ*) and pSTD343 (*lacI*) were transformed with either pSTD1060 (vector), pYK186 (pFecR), pYK188 (p3xFLAG-FecR), or pYK238 (p3xFLAG-FecR-PA). Cells were grown at 30 °C in M9-based medium containing 1 mM IPTG and 0.1 μM $FeCl_3$, with or without 1 mM $Na_3$-citrate, until mid-log phase. LacZ activities were measured, and relative activities normalized to that of the cells harboring pYK186 grown in the same medium with 1 mM $Na_3$-citrate. Data represent the means ± SD from two biologically independent experiments. Student's *t*-test was carried out to compare the values between the groups. *$P < 0.05$; **$P < 0.01$; ***$P < 0.001$; ns, $P > 0.05$.

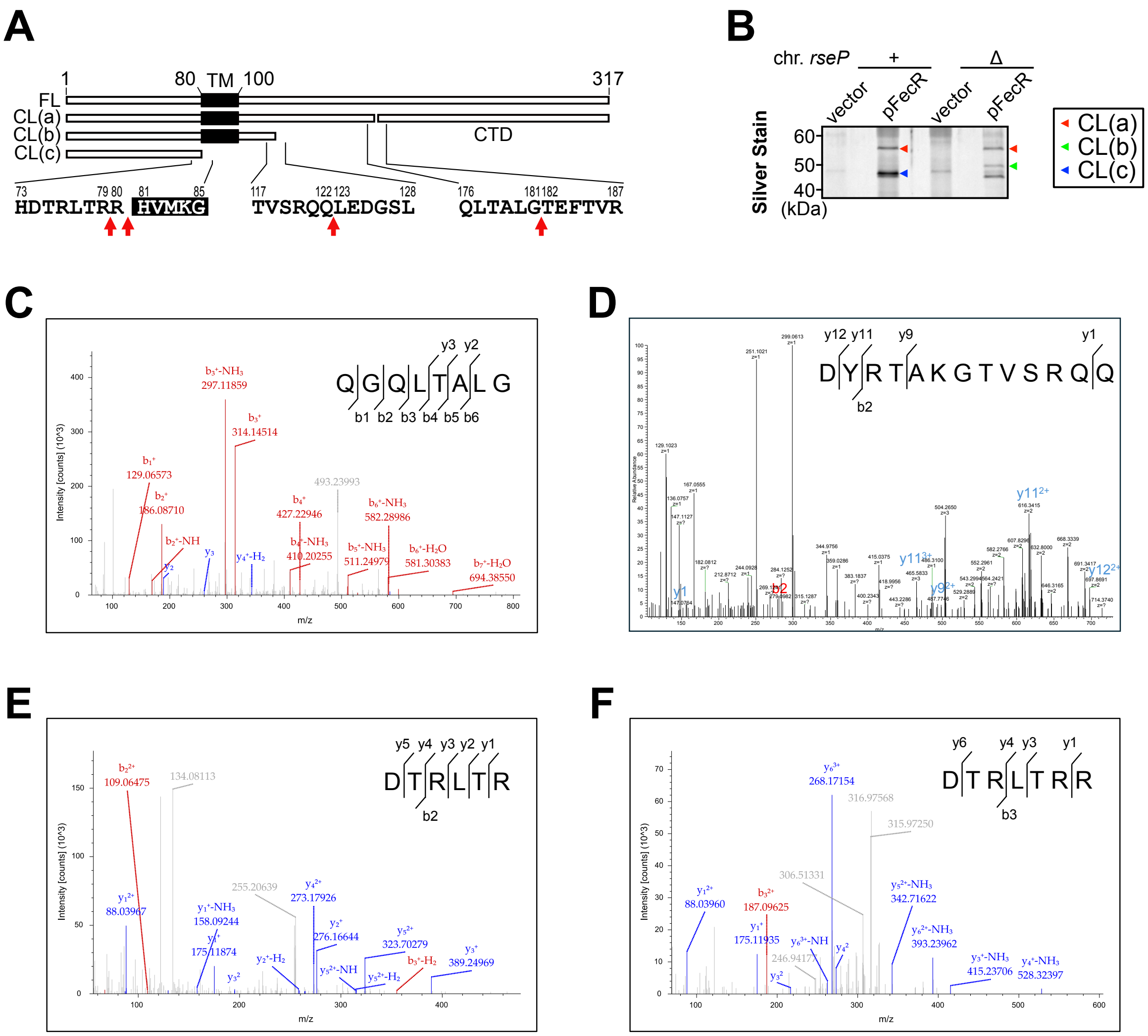


**Fig. S2. Identification of the C-terminal residues of the CL(a), CL(b), and CL(c) fragments.**

(A) Schematic representation of the FecR cleavage sites. The predicted transmembrane region is depicted as the black box. Cleavage sites identified by mass spectrometry (MS) analysis are indicated by red arrows. (B) Purification of the N-terminal cleavage fragments of FecR. YK167 (*rseP*+) or YK191 (Δ*rseP*) cells harboring either pTWV228 (vector), or pYK202 ($His_{10}$-MBP-FecR) were grown at 30 °C in M9-based medium containing 1 mM IPTG, 10 µM $FeCl_3$, and 1 mM $Na_3$-citrate until mid-log phase. Total cellular proteins were acid-precipitated, subjected to pull-down with TALON metal affinity resin (Takara Bio), and visualized by SDS-PAGE followed by silver staining. (C-F) MS/MS spectra of peptides containing the C-terminal residues from CL(a), CL(b), and CL(c). The C-terminal peptides containing the cleavage sites were identified using Sequest HT or manually. (C) The C-terminal peptide QGQLTALG (m/z = 787.4318, z = 1+, Xcorr = 1.99) was identified from the trypsin-digested peptide of the CL(a) fragment. (D) The C-terminal peptide DYRTAKGTVSRQQ (observed m/z = 503.9264, theoretical m/z = 503.9325, z = 3+, delta = 8.5ppm) was manually identified from the Asp-N-digested peptide of the CL(b) fragment. (E and F) The C-terminal peptides DTRLTR (m/z = 254.4803, z = 3+, Xcorr = 1.54) and DTRLTRR (m/z = 306.5138, z = 3+, Xcorr = 1.06) were identified from the Asp-N-digested peptides of the CL(c) fragment.

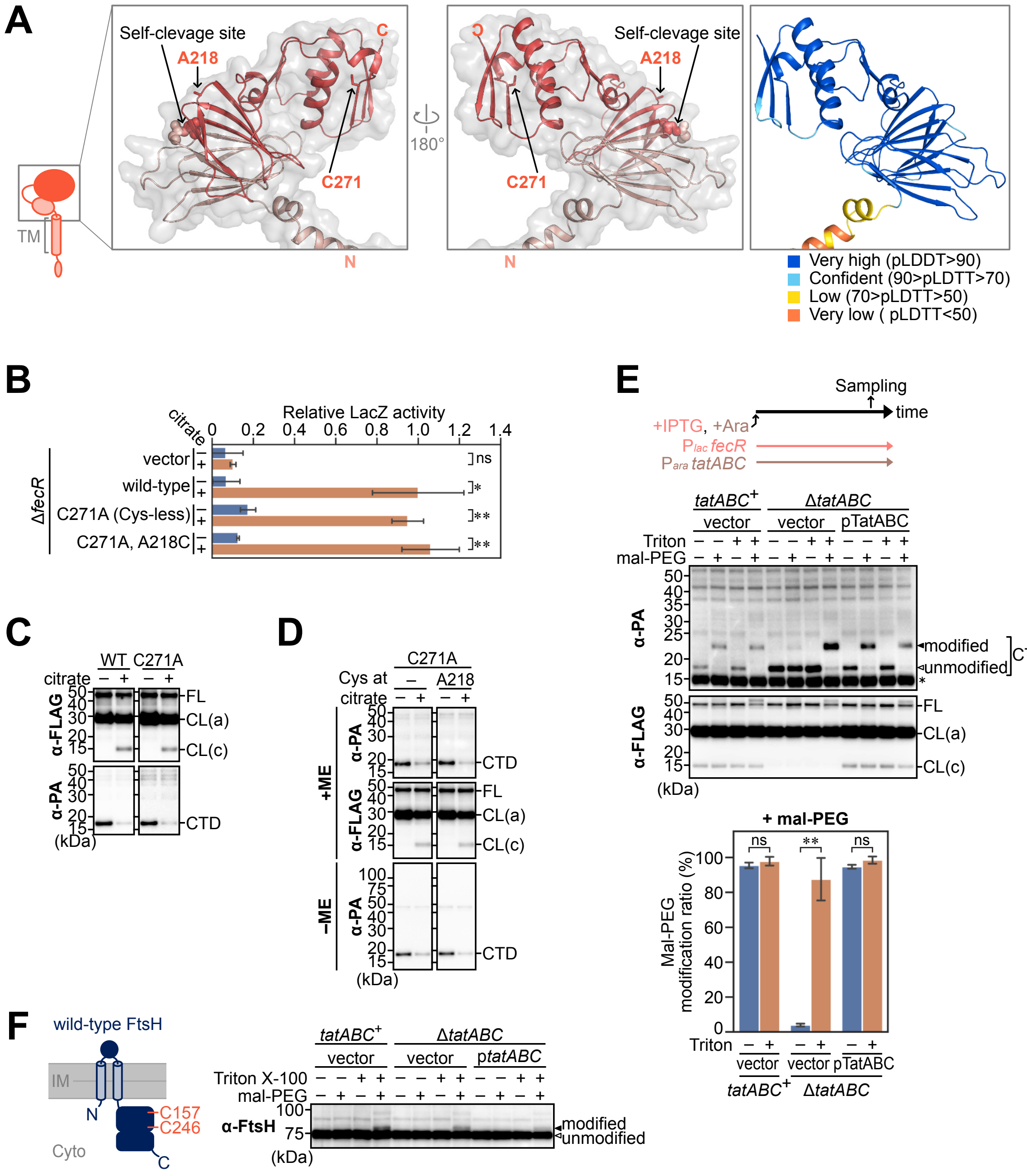


**Fig. S3. Mechanistic analysis of FecR membrane translocation.**

(A) AlphaFold3-predicted structure of full-length FecR. The regions corresponding to the FecR CL(a) and the CTD fragments are shown in pink and red, respectively. The C-terminal residue of CL(a) (G181) and the N-terminal residue of the CTD fragment (T182) are depicted as sphere models. The side chains of A218 and C271 are presented as stick models (A, left and middle). The per-residue model confidence score (pLDDT; predicted local distance difference test) values for the model structures are shown (A, right). (B) Ability of 3xFLAG-FecR-PA derivatives to transmit the ferric citrate signal. YK627 (Δ*fecR*) cells harboring pYK149 ($P_{fecA}$-*lacZ*) and pSTD343 (*lacI*) were transformed with either pSTD1060 (vector), pYK238 (3xFLAG-FecR-PA, wild-type), pYK1006 (3xFLAG-FecR-PA C271A, Cys-less), or pYK1102 (3xFLAG-FecR-PA C271A/A218C). Cells were grown at 30 °C in M9-based medium containing 1 mM IPTG and 0.1 μM $FeCl_3$, with or without 1 mM $Na_3$-citrate, until mid-log phase. LacZ activities were measured and normalized to that of the cells harboring pYK238 grown in the same medium with 1 mM $Na_3$-citrate. Data represent the means ± SD from two biologically independent experiments. Student's *t*-test was carried out to compare the values between the groups. *$P < 0.05$; **$P < 0.01$; ns, $P > 0.05$. (C and D) Cleavage profiles of 3xFLAG-FecR-PA derivatives in response to $Na_3$-citrate. (C) YK167 cells harboring pYK212 (3xFLAG-FecR-PA, WT) or pYK1001 (3xFLAG-FecR-PA C271A) and (D) YK167 cells harboring pYK1001 (3xFLAG-FecR-PA C271A) or pYK1025 (3xFLAG-FecR-PA C271A/A218C) were grown at 30 °C in M9-based medium containing 1 mM IPTG and 10 μM $FeCl_3$, with or without 1 mM $Na_3$-citrate (citrate), until mid-log phase. Total cellular proteins were acid-precipitated, dissolved in SDS sample buffer containing either 10% 2-mercaptoethanol (ME) (C) or 25 mM N-ethylmaleimide (NEM; to block free thiol groups) with or without 10% ME (D). Samples were analyzed by SDS-PAGE followed by immunoblotting with the indicated antibodies. FL, CL(a), CL(c), and CTD indicate the full-length protein, the N-terminal cleavage product CL(a), CL(c), and the C-terminal cleavage product CTD fragment, respectively. (E) TatABC-dependence of the cellular localization of the CTD fragment as assessed by the mal-PEG-2k modification assay. HM1742 (*tatABC*$^+$) or YK2238 (Δ*tatABC*) cells harboring pYK1025 (3xFLAG-FecR-PA C271A/A218C) were transformed with either pBAD33 (vector) or pYK1000 (pTatABC). A schematic workflow of the experiment is shown (E, upper). Cells were grown at 30 °C in M9-based medium containing 1 mM IPTG, 0.2 % arabinose, and 10 μM $FeCl_3$. Spheroplasts were prepared from these cells and treated with 2 mM mal-PEG-2k (mal-PEG) in the presence or absence of 1% Triton X-100 (Triton). Total cellular proteins were analyzed as described in (C) (E, middle). 'Modified' and 'unmodified' refer to mal-PEG-2k-modified and unmodified CTD fragments, respectively. An asterisk indicates lysozyme. The levels of mal-PEG-2k-modification of the CTD fragment were quantified and presented as a percentage of the total levels (mal-PEG-2k modified plus unmodified). Data represent the means ± SD from two biologically independent experiments. Student's *t*-test was carried out to compare the values between the groups. **$P < 0.01$; ns, $P > 0.05$ (E, lower). (F) Effect of TatABC deletion and overexpression on the mal-PEG-2k modifiability of the membrane-bound protease FtsH. A schematic representation of FtsH is shown (F, left). Wild-type FtsH contains two cysteine residues in its cytoplasmic domain. IM and Cyto indicate the inner membrane and cytoplasm, respectively. Cells were grown and treated with mal-PEG-2k as described in (E), and samples were analyzed by SDS-PAGE followed by immunoblotting with an anti-FtsH (α-FtsH) antibody (F, right).

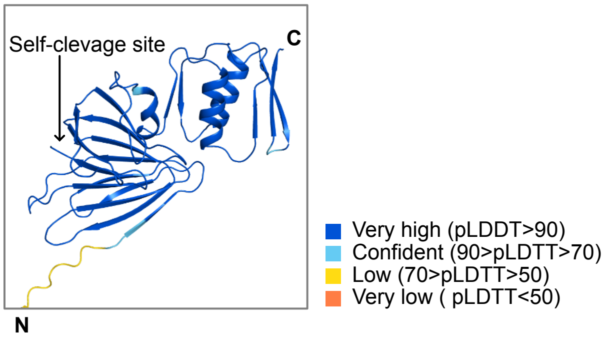


**Fig. S4. AlphaFold predicted structure of the FecR CL(a)-CTD fragment complex.**
AlphaFold3-predicted structure of the FecR CL(a)-CTD fragment complex with the per-residue pLDDT values is shown. This is the same model as shown in Fig. 4A.

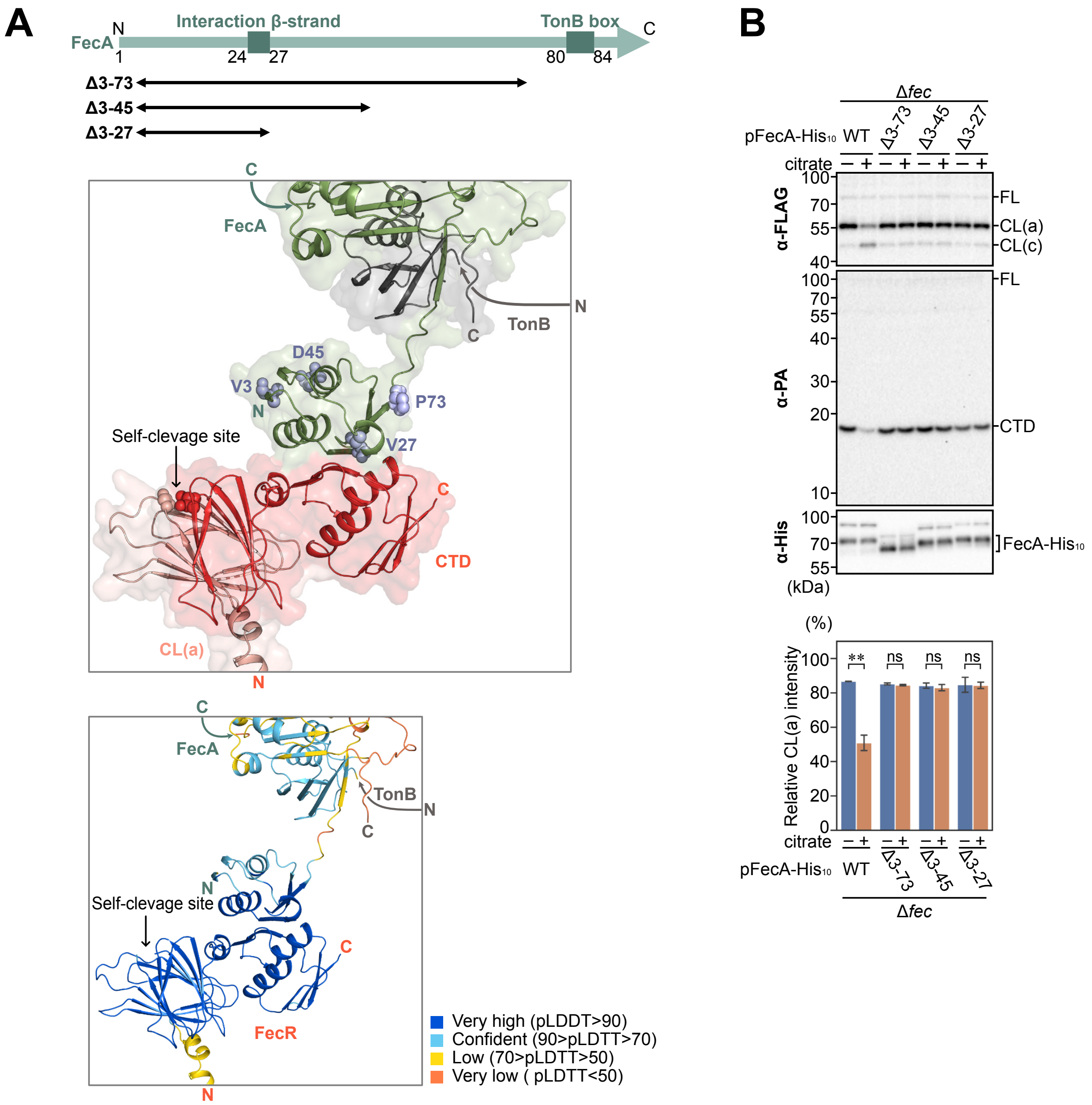


**Fig. S5. Effect of truncations in the FecA N-terminal region predicted to interact with the FecR CTD on signal-dependent cleavage of FecR.**

(A) AlphaFold3-predicted structure of the FecR CL(a)-CTD-FecA-TonB supercomplex. This is the same model as shown in Fig. 5A. A schematic representation of the N-terminal domain of FecA is shown (A, upper). The 'Interaction β-strand' in FecA, which is predicted to interact with the FecR CTD fragment, and the TonB box are indicated as green boxes. The structural model is depicted (A, middle and lower). FecR CL(a), the CTD fragment, FecA, and TonB are shown in pink, red, green, and gray, respectively (A, middle). Residues V3, V27, D45, and P73 of FecA are displayed as sphere models. The per-residue pLDDT values of the model structures are shown (A, lower). (B) Effect of FecA truncations on the FecR cleavage profile. YK167 ($fec^{+}$) or YK1602 (Δ*fec*) cells harboring pYK214 (3xFLAG-MBP-FecR-PA) were transformed with plasmids encoding the indicated FecA-$His_{10}$ derivatives. Cells were grown and total cellular proteins were analyzed as described in *SI Appendix*, Fig. S3C. The immunoblot image is shown (B, upper). Band intensities of fragments derived from 3xFLAG-MBP-FecR-PA in the indicated strains were quantified and the band intensities of CL(a) are presented as a percentage of the total intensities of all fragments.

Data represent the means ± SD from two biologically independent experiments. Student's *t*-test was carried out to compare the values between the groups. $^{**}P < 0.01$; ns, $P > 0.05$ (B, lower).

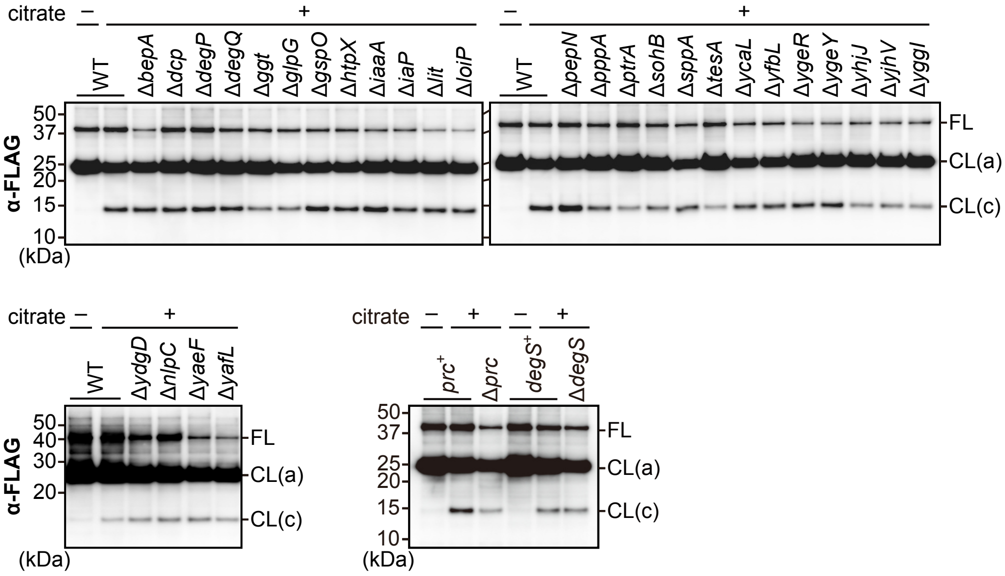


**Fig. S6. Screening for the protease(s) responsible for FecR CL(a) cleavage.**
Effect of the deletion of genes encoding membrane- or periplasm-localized peptidases on the FecR cleavage profile. BW25113-based strains lacking the indicated genes were transformed with pYK212 (3xFLAG-FecR-PA) and pSTD343 (*lacI*) (upper and lower left). Additionally, SN140 (*prc*$^+$), AD2215 (Δ*prc*), KK379 (*degS*$^+$), and KK372 (Δ*degS*) cells were transformed with pYK212 (lower right). Cells were grown and total cellular proteins were analyzed as described in *SI Appendix*, Fig. S3C.

**Table S1.** The strains, plasmids, and oligonucleotides used in this study.

| ***E. coli* strains** | | |
|---|---|---|
| Name | Genotype | Reference |
| MC4100 | F$^{-}$, *araD139* Δ*(argF-lac)U169 rpsL150 relA1 flbB5301 deoC1 ptsF25 rbsR* | (7) |
| YK627 | MC4100, Δ*fecR1* | (3) |
| CU141 | MC4100 /F' *lacI*$^{q}$ *Z*$^{+}$ *Y*$^{+}$ | (8) |
| HM1742 | CU141, *ara*$^{+}$ | (9) |
| YK167 | HM1742, Δ*rseA* | (3) |
| YK191 | HM1742, Δ*rseA* Δ*rseP*::*kan* | (3) |
| YK1196 | HM1742, Δ*rseA* Δ*tatABC* | This study |
| YK1602 | HM1742, Δ*rseA* Δ*fecIRABCDE* | This study |
| YK2234 | HM1742, Δ*rseA* Δ*tonB*::*kan* | This study |
| YK2238 | HM1742, Δ*tatABC*::*kan* | This study |
| BW25113 | F$^{-}$, *rrnB* Δ*lacZ4787 hsdR514* Δ(*araBAD*)*567* Δ(*rhaBAD*)*568 rph-1* | (5) |
| YK1131 | BW25113, Δ*tatABC*::*kan* | This study |
| YK1129 | BW25113, Δ*fecIRABCDE*::*kan* | This study |
| JW2556 | BW25113, Δ*rseA*::*kan,* KEIO collection | (10) |
| JW5195 | BW25113, Δ*tonB*::*kan,* KEIO collection | (10) |
| AD16 | Δ*pro-lac thi* /F' *lacI*$^{q}$ ZΔM15 *Y*$^{+}$ *pro*$^{+}$ | (11) |
| AD1840 | AD16, Δ*rseA*::*cat* Δ*rseP*::*kan* Δ*degS*::*tet* | (6) |
| JW2479 | BW25113, Δ*bepA*::*kan,* KEIO collection | (10) |
| JW1531 | BW25113, Δ*dcp*::*kan,* KEIO collection | (10) |
| JW0157 | BW25113, Δ*degP*::*kan,* KEIO collection | (10) |
| JW3203 | BW25113, Δ*degQ*::*kan,* KEIO collection | (10) |
| JW3412 | BW25113, Δ*ggt*::*kan,* KEIO collection | (10) |
| JW5687 | BW25113, Δ*glpG*::*kan,* KEIO collection | (10) |
| JW3297 | BW25113, Δ*gspO*::*kan,* KEIO collection | (10) |
| JW1818 | BW25113, Δ*htpX*::*kan,* KEIO collection | (10) |
| JW0812 | BW25113, Δ*iaaA*::*kan,* KEIO collection | (10) |
| JW2723 | BW25113, Δ*iaP*::*kan,* KEIO collection | (10) |
| JW1125 | BW25113, Δ*lit*::*kan,* KEIO collection | (10) |
| JW2903 | BW25113, Δ*loiP*::*kan,* KEIO collection | (10) |
| JW0915 | BW25113, Δ*pepN*::*kan,* KEIO collection | (10) |
| JW2939 | BW25113, Δ*pppA*::*kan,* KEIO collection | (10) |
| JW2789 | BW25113, Δ*ptrA*::*kan,* KEIO collection | (10) |
| JW1264 | BW25113, Δ*sohB*::*kan,* KEIO collection | (10) |
| JW1755 | BW25113, Δ*sppA*::*kan,* KEIO collection | (10) |
| JW0483 | BW25113, Δ*tesA*::*kan,* KEIO collection | (10) |
| JW0892 | BW25113, Δ*ycaL*::*kan,* KEIO collection | (10) |
| JW2266 | BW25113, Δ*yfbL*::*kan,* KEIO collection | (10) |
| JW2833 | BW25113, Δ*ygeR*::*kan,* KEIO collection | (10) |
| JW2840 | BW25113, Δ*ygeY*::*kan,* KEIO collection | (10) |
| JW3495 | BW25113, Δ*yhjJ*::*kan,* KEIO collection | (10) |
| JW4246 | BW25113, Δ*yjhV*::*kan,* KEIO collection | (10) |
| JW2911 | BW25113, Δ*yggI*::*kan,* KEIO collection | (10) |
| JW1590 | BW25113, Δ*ydgD*::*kan,* KEIO collection | (10) |
| JW1698 | BW25113, Δ*nlpC*::*kan,* KEIO collection | (10) |
| JW5016 | BW25113, Δ*yaeF*::*kan,* KEIO collection | (10) |
| JW0217 | BW25113, Δ*yafL*::*kan,* KEIO collection | (10) |
| SN140 | CU141, *ompT*::*kan* | (12) |
| AD2215 | SN140, *prc*::*cat* | laboratory stock |
| KK379 | CU141, Δ*rseA*::*cat* | A gift of K. Kanehara |
| KK372 | KK379, Δ*degS*::*tet* | (13) |

| **Plasmids** | | | |
|---|---|---|---|
| Name | Vector | Encoded proteins or descriptions | Reference |
| pTWV228 | | pBR322-based vector; $P_{lac}$, $Amp^R$ | Takara Bio |
| pUC118 | | pBR322-based vector; $P_{lac}$, $Amp^R$ | Takara Bio |
| pSTD689 | | pACYC184-based vector; $P_{lac}$, $Spc^R$ | (14) |
| pSTD689c | | pSTD689-based constitutive expression vector | (15) |
| pBAD33 | | pACYC184-based vector; $P_{ara}$, $Cm^R$ | (16) |
| pMW118 | | pSC101-based vector; $P_{lac}$, $Amp^R$ | Nippon Gene |
| pSTV29 | | pACYC184-based vector; $P_{lac}$, $Cm^R$ | Takara Bio |
| pSTD1060 | | pBR322-based vector; $P_{lac}$, $Spc^R$ | (3) |
| pKD13 | | Template for the PCR of cassette construction | (5) |
| pKD46 | | λ-Red recombinase system | (5) |
| pCP20 | | FLP recombinase | (17) |
| pYK186 | pSTD1060 | FecR | (3) |
| pYK130 | pTWV228 | 3xFLAG-FecR | (3) |
| pYK188 | pSTD1060 | 3xFLAG-FecR | (3) |
| pYK336 | pTWV228 | 3xFLAG-FecR CL(a)-mimic (T182stop) | This study |
| pYK341 | pSTD1060 | 3xFLAG-FecR CL(a)-mimic (T182stop) | This study |
| pYK212 | pTWV228 | 3xFLAG-FecR-PA | This study |
| pYK238 | pSTD1060 | 3xFLAG-FecR-PA | This study |
| pYK1001 | pTWV228 | 3xFLAG-FecR (C271A)-PA | This study |
| pYK1006 | pSTD1060 | 3xFLAG-FecR (C271A)-PA | This study |
| pYK1025 | pTWV228 | 3xFLAG-FecR (C271A, A218C)-PA | This study |
| pYK1102 | pSTD1060 | 3xFLAG-FecR (C271A, A218C)-PA | This study |
| pYK1048 | pTWV228 | 3xFLAG-FecR (C271A, K115C)-PA | This study |
| pYK1060 | pTWV228 | 3xFLAG-FecR (C271A, N266C)-PA | This study |
| pYK1079 | pTWV228 | 3xFLAG-FecR (C271A, K115C, N266C)-PA | This study |
| pYK147 | pTWV228 | 3xFLAG-MBP-FecR | (3) |
| pYK172 | pTWV228 | 3xFLAG-MBP-FecR CL(a)-mimic (T182stop) | (3) |
| pYK214 | pTWV228 | 3xFLAG-MBP-FecR-PA | This study |
| pYK2004 | pTWV228 | 3xFLAG-MBP-FecR (C271A)-PA | This study |
| pYK2006 | pTWV228 | 3xFLAG-MBP-FecR (C271A, K115C)-PA | This study |
| pYK2008 | pTWV228 | 3xFLAG-MBP-FecR (C271A, N266C)-PA | This study |
| pYK2010 | pTWV228 | 3xFLAG-MBP-FecR (C271A, K115C, N266C)-PA | This study |
| pYK202 | pTWV228 | $His_{10}$-MBP-FecR | This study |
| pYK364 | pTWV228 | FecA | This study |
| pYK367 | pSTD689 | FecA | This study |
| pYK369 | pSTD689c | FecA | This study |
| pYK2023 | pSTD689 | FecA-$His_{10}$ | This study |
| pYK2026 | pSTD689 | FecA (T24P)-$His_{10}$ | This study |
| pYK2027 | pSTD689 | FecA (L25P)-$His_{10}$ | This study |
| pYK2028 | pSTD689 | FecA (S26P)-$His_{10}$ | This study |
| pYK2029 | pSTD689 | FecA (V27P)-$His_{10}$ | This study |
| pYK2030 | pSTD689 | FecA (Δ3-73)-$His_{10}$ | This study |
| pYK2031 | pSTD689 | FecA (Δ3-45)-$His_{10}$ | This study |
| pYK2032 | pSTD689 | FecA (Δ3-27)-$His_{10}$ | This study |

| | | | |
|---|---|---|---|
| pYK301 | pTWV228 | TatA, TatB, TatC | This study |
| pYK345 | pSTD689 | TatA, TatB, TatC | This study |
| pYK1000 | pBAD33 | TatA, TatB, TatC | This study |
| pYK1100 | pSTD689 | TonB | This study |
| pYK149 | pMW118 | $P_{fecA}$-*lacZ* | (3) |
| pSTD343 | pSTV29 | LacI | (18) |

**Oligonucleotides**

| No. | Name | Sequence |
|---|---|---|
| P1 | tatABC_disrupt(+) | GCGGCTTTGTTTAATCATCATCTACCACAGAGGAACATGTGTGTAGGCTGGAGCTGCTTC |
| P2 | tatABC_disrupt(-) | CTGTACTCCATATGACAACCGCCCTGACGGGCGGTTGAATATTCCGGGGATCCGTCGACC |
| P3 | fec_disrupt(+) | TAATATGACTACGTGATAATTAACTTTTGATGCACTCCGCGTGTAGGCTGGAGCTGCTTC |
| P4 | fec_disrupt(-) | TTTCATTCAGTCGTGGTTTGGTTCTTACGGCCTGTGCAATATTCCGGGGATCCGTCGACC |
| P11 | PA_oligo(+) | AACATTTCACCACTGGTCGACGGCGTTGCCATGCCAGGTGCCGAAGATGATGTGGTGTAAAAGCTTGGCACT |
| P12 | PA_oligo(-) | AGTGCCAAGCTTTTACACCACATCATCTTCGGCACCTGGCATGGCAACGCCGTCGACCAGTGGTGAAATGTT |
| P13 | PA_insert_vec(+) | TAAAAGCTTGGCACTGGCCG |
| P14 | PA_insert_vec(-) | CAGTGGTGAAATGTTTATCCAG |
| P15 | BamH1-FecRTM(+) | TATCACCAAGGGATCCGATACCCGCCTCACCCGC |
| P16 | FecR-Sal1(-) | TGGCAACGCCGTCGACCAGTGGTGAAATGTTTATCCAGTAC |
| P17 | SacI-His-KpnI(+) | CGGTACAAGGAGGAAGAGCAAATGCATCACCATCACCATCACCATCACCATCACTCGGTAC |
| P18 | SacI-His-KpnI(-) | CGAGTGATGGTGATGGTGATGGTGATGGTGATGCATTTGCTCTTCCTCCTTGTACCGAGCT |
| P19 | EcoRI-SD-FecA(+) | CGGAATTCGAGCTCAAGGAGGAAGAGCAAATGACGCCGTTACGCGTT |
| P20 | FecA-HindIII(-) | CCCAAGCTTTCAGAACTTCAACGACCCC |
| P21 | FecA-GSGS-His(+) | CACCACCATCACCATCATCATCACCATCACTGAAAGCTTGGCACTGG |
| P22 | FecA-GSGS-His(-) | GATGGTGATGGTGGTGGCTGCCGCTACCGAACTTCAACGACCCCTG |
| P23 | FecA_Del3-73(+) | CTGCACAGGCGCCCGCACCAAAAGAA |
| P24 | FecA_Del3-73(-) | CGGGCGCCTGTGCAGCAAAAGCGGA |
| P25 | FecA_Del3-45(+) | CTGCACAGGTCGAGAGCGGCCTGCAAC |
| P26 | FecA_Del3-45(-) | TCTCGACCTGTGCAGCAAAAGCGGAAAAC |
| P27 | FecA_Del3-27(+) | CTGCACAGGACGCCAGCCTGACGCGC |
| P28 | FecA_Del3-27(-) | TGGCGTCCTGTGCAGCAAAAGCGGAAAACG |
| P29 | KpnI-tatABC(+) | CGGGGTACCTAATCATCATCTACCACAGAG |
| P30 | tatABC-SalI(-) | ACGCGTCGACTTATTCTTCAGTTTTTTCGCTTTC |
| P31 | noTag_TonB(+) | AAGGAGGAAGAGCAAATGACCCTTGATTTACCTCGCCG |

| | | |
|---|---|---|
| P32 | noTag_TonB(-) | GCCAGTGCCAAGCTTTTACTGAATTTCGGTGGTGCC |
| P33 | vec_pYK367(+) | AAGCTTGGCACTGGCCGTCG |
| P34 | vec_pYK367(-) | TTGCTCTTCCTCCTTGAGCTCG |
| P101 | PURE_F-FecR-PA(+) | AAGGAGATATACCAATGGACTACAAAGACCATGACG |
| P102 | PURE_F-FecR-PA(-) | GGATTAGTTATTCATTACACCACATCATCTTCGGC |
| P103 | T7PRO-SD(+) | GAAATTAATACGACTCACTATAGGGAGACCACAACGGTTTCCCTCTAGAAATAATTTTGTTTAACTTTAAGAAGGAGATATACCA |

**Dataset S1 (separate file).** The predicted structural model of FecR CL(a)/CTD fragment complex.

**Dataset S2 (separate file).** The predicted structural model of the FecR CL(a)/CTD fragment/FecA/ TonB supercomplex.

**Dataset S3 (separate file).** The predicted structural model of the full-length form of FecR.
